# *In vivo* direct imaging of neuronal activity at high temporo-spatial resolution

**DOI:** 10.1101/2021.05.21.444581

**Authors:** Phan Tan Toi, Hyun Jae Jang, Kyeongseon Min, Sung-Phil Kim, Seung-Kyun Lee, Jongho Lee, Jeehyun Kwag, Jang-Yeon Park

## Abstract

There has been a longstanding demand for noninvasive neuroimaging methods capable of detecting neuronal activity at both high temporal and spatial resolution. Here, we propose a novel method that enables Direct Imaging of Neuronal Activity for functional MRI (termed DIANA-fMRI) that can dynamically image spiking activity in milliseconds precision, while retaining the original benefit of high spatial resolution of MRI. DIANA-fMRI was demonstrated through *in vivo* mice brain imaging at 9.4 T applying electrical whisker-pad stimulation, directly imaging the spiking activity as well as capturing its sequential propagation along the thalamocortical pathway, as further confirmed through *in vivo* spike recording and optogenetics. DIANA-fMRI will open up new avenues in brain science by providing a deeper understanding of the brain’s functional organization including neural networks.

## Introduction

Advanced noninvasive neuroimaging methods provide valuable information on the brain’s functional organization, but they have obvious pros and cons in terms of temporal and spatial resolution. Functional magnetic resonance imaging (fMRI) using blood-oxygenation-level-dependent (BOLD) effect provides good spatial resolution in the order of millimeters, but has a poor temporal resolution in the order of seconds due to slow hemodynamic responses to neuronal activation (*1*), providing only indirect information on neuronal activity through neurovascular coupling. In contrast, electroencephalography (EEG) and magnetoencephalography (MEG) provide excellent temporal resolution in the millisecond range, but spatial information is limited to centimeter scales (*2*). Thus, while harnessing the high spatial resolution of MRI, enhancement of MRI-based temporal resolution up to that of EEG or MEG that can directly measure neuronal activity in the order of milliseconds is imperative to advance the understanding of *in vivo* brain, especially to elucidate the causal link between the *in vivo* neuronal activities and brain function.

Over the past decades, many attempts have been made in exploring the possibility of using MRI to directly image neuronal activity (*3*). Most of the studies were based on the neuronal current models, where neuronal currents flowing along the axon produce an ultraweak circumferential magnetic field on the order of nanotesla (nT) (*4, 5*), thereby locally changing the phase and magnitude of MR signals. Phantom studies demonstrated small phase shifts (< 1°) induced by magnetic field changes ≤ 1 nT when injecting electrical currents through wires in a gel phantom (*4, 6–8*). The *in vitro* studies using hemoglobin-free biological objects such as rat neuron cells (*9*), snail ganglia (*10, 11*), isolated brain of turtle (*8, 12*), and octopus (*13*) also reported changes in the magnitude of MR signals (e.g., 0.01 – 5.5%) induced by the applied stimulus. More interestingly, some *in vivo* studies have argued that they succeeded in directly detecting the human brain activation *in vivo* in response to event-related tasks (*14–19*), but they have not been replicated in later attempts (*20–24*).

Despite many efforts to use MRI to directly detect neuronal activity, no firm consensus has been reached on its feasibility yet, especially in *in vivo* studies. Here, we propose a novel method that enables Direct Imaging of Neuronal Activity for functional MRI (termed DIANA-fMRI) by increasing the temporal resolution up to the temporal precision of neuronal activity in milliseconds, while retaining the original benefit of high spatial resolution of MRI.

## Results

### Magnetic resonance imaging of neuronal activity at milliseconds temporal resolution

To implement high temporal resolution in milliseconds scale, we combined the line scanning method (*25*) and fast low angle shot (FLASH) gradient-echo imaging, where a single line of *k*-space was repeatedly acquired during each interstimulus interval, and different *k*-space lines were acquired in different periods. In this scheme, each stimulation period adds one line of the *k*-space to all the time-series images within the period (Fig. 1A) (*26, 27*), and the repetition time (TR) of the FLASH sequence exactly determines the temporal resolution of the dynamic imaging in DIANA-fMRI. Under this paradigm, electrical stimulation was repeatedly applied at 200 ms interstimulus intervals, determined by 5 ms TR multiplied by 40 frames in the time series. DIANA-fMRI showed similar performance to regular two-dimensional FLASH imaging in terms of signal-to-noise ratio (SNR) and temporal SNR (tSNR) (fig. S1). To test whether DIANA-fMRI can directly detect neuronal activity, we delivered electrical stimulation to the left whisker-pad (strength, 0.5 mA; duration, 0.5 ms; frequency, 5 Hz) of anesthetized mice placed inside the 9.4 T scanner and imaged a single 1 mm coronal brain slice which included the right barrel field of primary somatosensory cortex (S1BF) that is contralateral to the left whisker pad (Fig. 1B). In response to the electrical whisker-pad stimulation, a statistically significant increase in the DIANA signal was observed from the contralateral S1BF compared to the pre-stimulus signal (0.157 ± 0.015%, *p* < 0.001, 5 mice) (Fig. 1, C to E; fig. S2), whereas there were no significant changes in the unstimulated control mice, postmortem mice (Fig. 1, C to E; fig. S2) nor in sham experiments performed using an agar phantom (fig. S3). Most interestingly, the peak DIANA signal occurred with a latency of 24.00 ± 2.92 ms after the whisker-pad stimulation onset (Fig. 1, C, D and F), indicating that DIANA can detect whisker-pad stimulation-evoked responses by achieving high temporal resolution in milliseconds range. To find the neural correlates of DIANA-fMRI *in vivo*, the same whisker-pad stimulation paradigm as in Fig. 1B was repeated but now in mice implanted with a 32-channel silicon probe in the S1BF (Fig. 1G) to record the local field potential (LFP) and single-unit spike activities (Fig. 1H), from which their latencies were analyzed (Fig. 1, I to K). The peak of whisker-pad stimulation-evoked LFP had a latency of 39.48 ± 1.84 ms (12 mice), which was significantly slower than the latency of DIANA response (*p* < 0.001, Fig. 1, I and K). However, the peak spike firing rates of whisker-pad stimulation-responsive single units had a latency of 26.44 ± 1.24 ms (27 units from 10 mice) in the post-stimulus time histogram (PSTH) (Fig. 1J), which, surprisingly, was similar to the DIANA response latency as their latencies were not statistically different (*p* > 0.05, Fig. 1K). Additionally, other temporal spike characteristics such as time-to-first spike latency as well as median and mode of whisker-pad stimulation-responsive spike timings were also similar to the DIANA response latency (fig. S4). DIANA response amplitudes and spike firing rates both increased with increasing strength of electrical whisker-pad stimulation with little change in the latencies of their peaks (fig. S5). Together, these results show for the first time that DIANA-fMRI can measure transient neuronal activity with milliseconds temporal resolution that is comparable to the single-unit spike recordings *in vivo*.

**Fig. 1.**
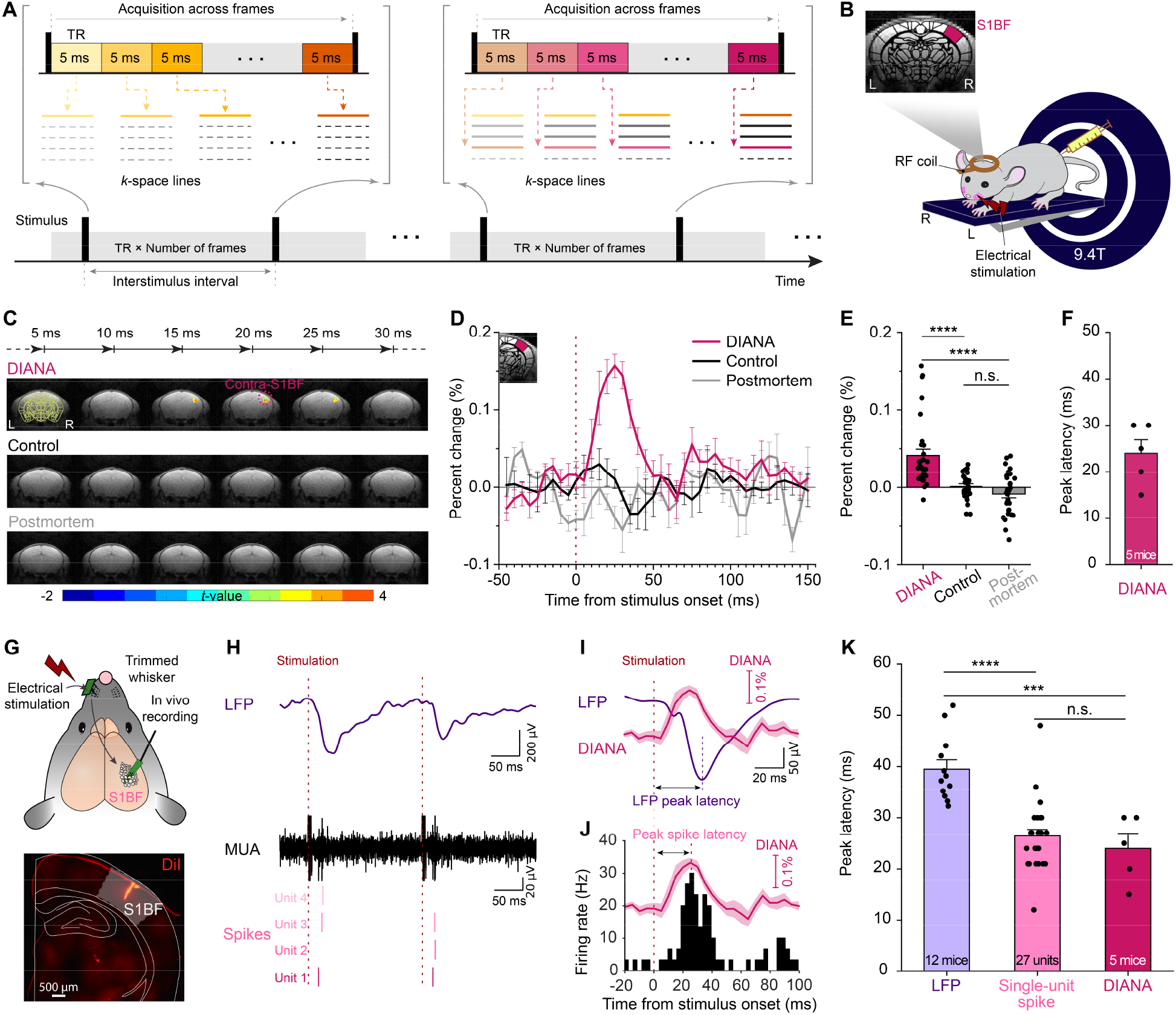
DIANA-fMRI: Direct Imaging of Neuronal Activity for functional MRI with high temporo-spatial resolution. (**A**) DIANA-fMRI acquisition scheme. (**B**) Illustration of DIANA-fMRI experiment to image S1BF applying electrical stimulation to left whisker pad in an anesthetized mouse on a 9.4 T scanner. (**C** to **E**) Time series of *t*-value maps of DIANA-fMRI in 5 ms temporal resolution (C), percent changes in DIANA signals (D) and mean signal changes during post-stimulation (E) with whisker-pad stimulation (magenta, *n* = 5 mice), without stimulation (control, black, *n* = 5 mice), and in postmortem condition (gray, *n* = 4 mice). (**F**) Latency of peak DIANA response. (**G**) Top: Illustration of electrophysiological recording in mice *in vivo* with a 32-channel silicon probe implanted in the contralateral S1BF applying electrical whisker-pad stimulation, Bottom: Electrode track marking using a fluorescent lipophilic dye (DiI). (**H**) A representative local field potential (LFP) signal (purple trace, top) and multi-unit activity (MUA) (black trace, middle) from which single-unit spikes (bottom) were analyzed. Spikes of each single unit are displayed in different shades of magenta. (**I** and **J**) LFP (I) and post-stimulus time histogram (PSTH) of the whisker-pad stimulation-responsive single units over time in the contralateral S1BF (J) with DIANA signals superimposed (magenta trace) for comparison. (**K**) Bar graph showing the latencies of LFP (purple, *n* = 12 mice), the peak spike firing rate of whiskerpad stimulation-responsive single units (light magenta, *n* = 27 units from 10 mice), and the DIANA response (magenta, *n* = 5 mice). Vertical dotted lines indicate the whisker-pad stimulation onset time (D, and H to J) and latency of either peak LFP (I) or peak spike firing rate (J). All data are mean ± SEM. ***: *p* < 0.001, ****: *p* < 0.0001, n.s.: *p* > 0.05 for one-way ANOVA with Bonferroni *post hoc* test.

### Temporo-spatial imaging of neuronal activity propagation

Since somatosensory stimulus-evoked spikes propagate to S1BF via thalamus in the thalamocortical pathway (*28, 29*), we next explored if the high temporo-spatial resolution (5 ms, 0.22 mm) of DIANA-fMRI can also capture the propagation of spikes. When a 1 mm coronal brain slice that included both the thalamus and S1BF was imaged applying electrical whisker-pad stimulation (Fig. 2A), while conventional BOLD-fMRI showed concomitant activation of thalamus, contra- and ipsilateral S1BF (Fig. 2B and fig. S6), DIANA-fMRI showed statistically significant DIANA response which was sequentially activated in the order of thalamus (0.157 ± 0.011%) and, almost simultaneously, contralateral S1BF (0.161 ± 0.009%) and ipsilateral S1BF (0.047 ± 0.014%) (Fig. 2, C and D; fig. S7, A and B) with latencies of 11.50 ± 0.76 ms, 25.00 ± 1.49 ms, and 25.00 ± 3.16 ms (Fig. 2E and fig. S7C) following stimulation onset, respectively. Further cross-correlation analysis of the time series confirmed that thalamic responses precede contra- and ipsilateral S1BF by 10 - 15 ms (fig. S8). To see if the temporo-spatial propagation of DIANA-fMRI matches that of spikes in the thalamocortical pathway, we performed simultaneous single-unit recordings in the thalamus and S1BF using two silicon probes with electrical whiskerpad stimulation (Fig. 2, F to I). Consistent with DIANA responses, latencies of peak spike firing rates of the stimulation-responsive single units occurred in the order of the thalamus (9.52 ± 0.90 ms, 23 units from 10 mice), contralateral S1BF (24.52 ± 1.43 ms, 23 units from 5 mice) (Fig. 2, I and J) and ipsilateral S1BF (30.00 ± 1.73 ms, 3 units from 1 mouse) (fig. S9). Such sequential propagation of spikes in the thalamus and the S1BF was statistically similar to those observed from DIANA responses (Fig. 2E and fig. S10). LFP also showed sequential propagation in the thalamus (31.78 ± 3.67 ms), contralateral S1BF (42.69 ± 3.04 ms), and ipsilateral S1BF (57.67 ± 5.33 ms), but they were significantly slower than the DIANA responses (fig. S10). These results demonstrate that DIANA-fMRI can capture spike propagation across functionally-defined thalamocortical pathway with high temporo-spatial resolution.

**Fig. 2.**
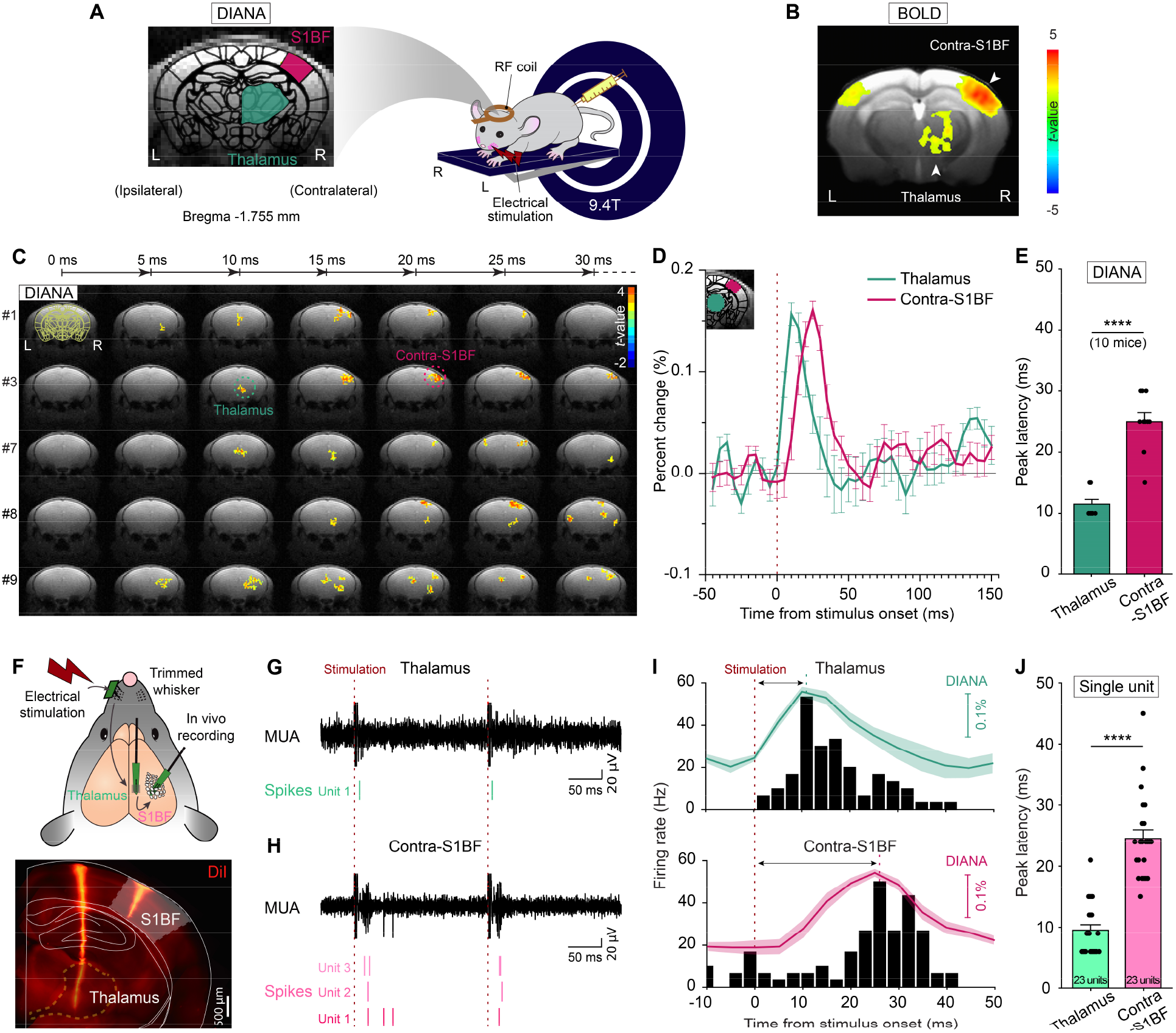
High temporo-spatial resolution of DIANA-fMRI captures thalamocortical spike propagation. (**A**) Illustration of DIANA-fMRI experiment to image contralateral S1BF and thalamus applying electrical stimulation to left whisker pad in an anesthetized mouse on a 9.4 T scanner (right) and brain imaging of a coronal slice containing both thalamus and S1BF regions (left). (**B**) BOLD activation map obtained as a reference (*n* = 10 mice). (**C** to **E**) Time series of *t*-value maps of DIANA-fMRI for 30 ms after whisker-pad stimulation in 5 ms temporal resolution from 5 representative mice (C), percent changes in DIANA signals (D), and bar graph showing the mean latencies of peak DIANA responses from thalamus (green) and contralateral S1BF (magenta) (E) (*n* = 10 mice, ****: *p* < 0.0001, paired *t*-test). (**F**) Top: Illustration of electrophysiological recording in mice *in vivo* with silicon probes implanted in the thalamus and the contralateral S1BF applying electrical whisker-pad stimulation, Bottom: Electrode track marking using a fluorescent lipophilic dye (DiI). (**G** and **H**) A representative multi-unit activity (MUA) (black trace, top) from which single-unit spikes (bottom) were analyzed in the thalamus (green) (G) and the contralateral S1BF (magenta) (H). (**I**) Post-stimulus time histogram (PSTH) of the whisker-pad stimulation-responsive single units over time in the thalamus (top) and contralateral S1BF (bottom) with DIANA signals superimposed for comparison. (**J**) Bar graph showing the latencies of peak spike firing rates of whisker-pad stimulation-responsive single units recorded from the thalamus (light green, *n* = 23 units from 10 mice) and contralateral S1BF (light magenta, *n* = 23 units from 5 mice). Vertical dotted lines indicate the whisker-pad stimulation onset time (red) (D, and G to I) and latency of peak spike firing rate (thalamus, green; contralateral S1BF, magenta) (I). (****:*p* < 0.0001, unpaired *t*-test). All data are mean ± SEM.

### Optogenetic DIANA-fMRI experiment

Although the temporal characteristics of DIANA responses and electrophysiologically recorded spikes *in vivo* are statistically similar, these measurements are somewhat limited since they were not measured simultaneously. To more directly verify that DIANA is capable of imaging spike activity *in vivo*, we employed an optogenetic fMRI scheme. DIANA responses were measured during optogenetic activation of Channelrhodopsin2 (ChR2)-expressing excitatory neurons in the S1BF with 473 nm blue light delivered through chronically implanted fiber-optic cannula (Fig. 3A). Immunostaining showed that stereotaxically injected AAV5-CamKIIa-hChR2(ET/TC)-mCherry (*30*) into S1BF expressed ChR2-mCherry in excitatory neurons across all cortical layers (Fig. 3B). During blue light stimulation (intensity, 50 mW/mm^2^; duration, 20 ms), DIANA responses were acquired as a time series of 50 images every 5 ms from a 1 mm coronal brain slice containing both the thalamus and S1BF (Fig. 3C). We found statistically significant DIANA signal change in the S1BF (0.200 ± 0.070%) with response latencies of 13.75 ± 4.27 ms, which was followed by significant DIANA signal change in the thalamus (0.217 ± 0.081%) 37.50 ± 8.78 ms after stimulation onset (Fig. 3, D and E), directly imaging optogenetic blue light stimulation-evoked neuronal activities as well as capturing the feedback spike propagation in the cortico-thalamic pathway (*29, 31, 32*). Indeed, when we recorded blue light stimulation-evoked single-unit activities from the S1BF and the thalamus of the same mice that were used in optogenetic DIANA-fMRI experiment (Fig. 3F), peak spike firing rate of neurons in S1BF occurred 9.06 ± 1.59 ms after stimulation onset (34 units from 8 mice), followed by thalamic peak spike firing rate occurring at 21.86 ± 2.76 ms (7 units from 4 mice) (Fig. 3, G to I), showing feedback spike propagation in the cortico-thalamic pathway. In addition, both DIANA responses and peak spike firing rates increased with increasing blue light stimulation duration (fig. S11), indicating similarities of DIANA responses and single-unit activities. Although feedback propagation of LFP in the cortico-thalamic pathway was also observed (S1BF, 22.34 ± 0.87 ms; thalamus, 77.55 ± 4.61 ms), peak LFP latencies were significantly slower than those of optogenetic DIANA responses (fig. S12). Together, optogenetic DIANA and corresponding *in vivo* single-unit recordings further confirm that the DIANA response is indeed consequences of direct detection of the spike activity *in vivo*.

**Fig. 3.**
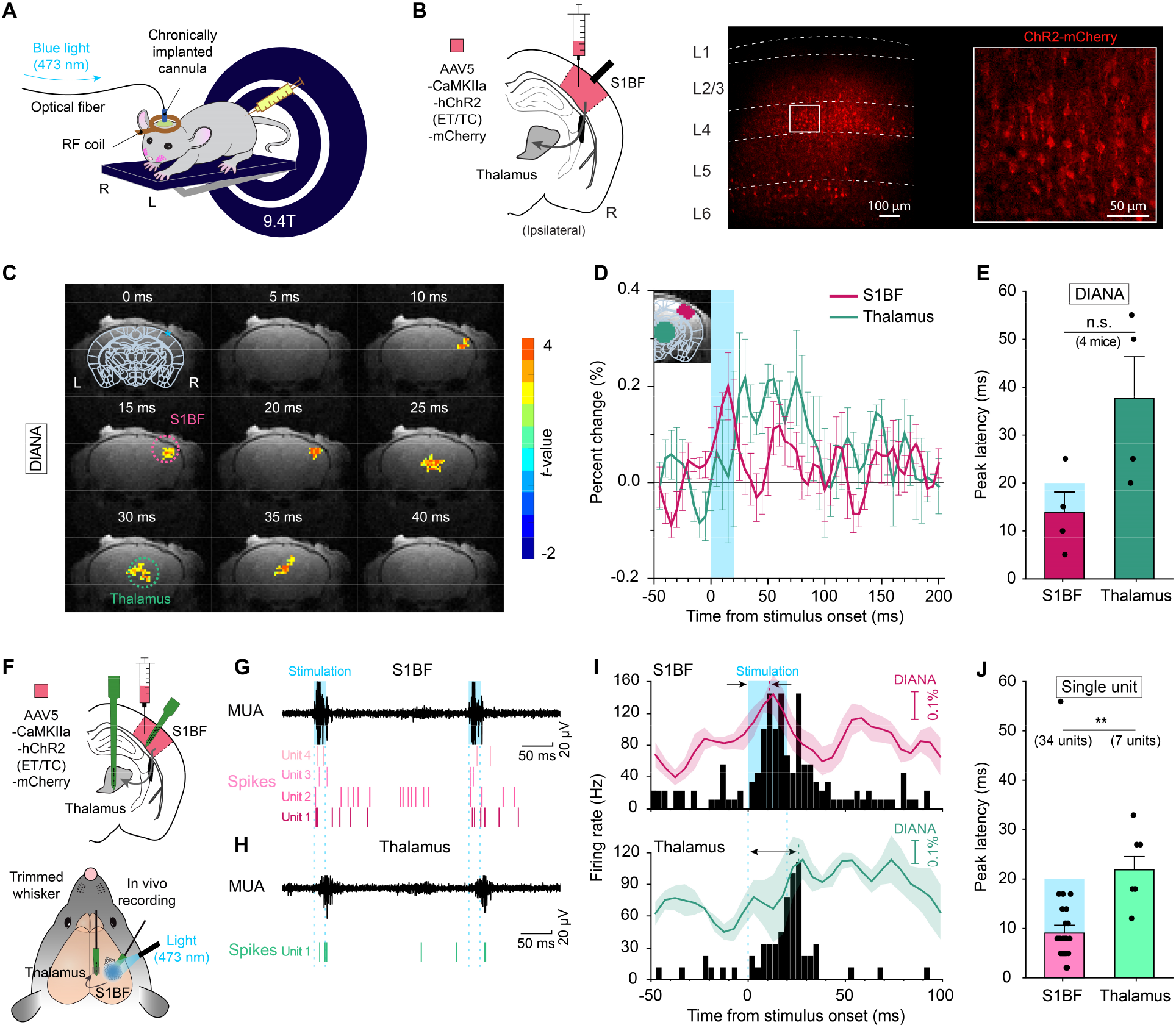
Optogenetic DIANA-fMRI: DIANA responses directly detect optogenetic stimulation-evoked spikes. (**A**) Illustration of optogenetic DIANA-fMRI where a fiber-optic cannula is implanted in the mouse S1BF for blue light (473 nm) stimulation. (**B**) Illustration of injection of AAV5-CaMKIIa-hChR2(ET/TC)-mCherry in S1BF (left) which expressed Channelrhodopsin (ChR2) to excitatory neurons across all layers of S1BF as confirmed by confocal imaging of mCherry-expressing excitatory neurons (right). (**C**) Time series of *t*-value maps of DIANA-fMRI from the S1BF and thalamus of a representative mouse (cluster size > 5 voxels) imaged for 40 ms after blue light (473 nm) stimulation onset (intensity, 50 mW/mm^2^; duration, 20 ms). (**D** and **E**) Percent changes in DIANA signals (D) and bar graph showing the mean latencies of peak DIANA responses in the contralateral S1BF (magenta) and thalamus (green) (E) (*n* = 4 mice, n.s.: *p* > 0.05, paired *t*-test). Blue shading indicates the period of blue light stimulation. (**F**) Illustration of simultaneous electrophysiological recordings *in vivo* in the thalamus and S1BF in mice injected with AAV5-CaMKIIa-hChR2(ET/TC)-mCherry in S1BF, using the same blue light stimulation as in optogenetic DIANA-fMRI. (**G** and **H**) A representative multi-unit activity (MUA) (black trace, top) from which single-unit spikes (bottom) were analyzed in the contralateral S1BF (G) and the thalamus (H). (**I**) Post-stimulus time histogram (PSTH) of the blue light stimulation-responsive single units over time in the S1BF (top) and thalamus (bottom) with DIANA signals superimposed for comparison. (**J**) Bar graph showing the latencies of peak spike firing rates following the blue light stimulation in the S1BF (*n* = 34 units from 8 mice) and thalamus (*n* = 7 units from 4 mice) (**: *p* < 0.01, unpaired *t*-test). Blue shading/dotted lines indicate the period of optogenetic stimulation in S1BF (D, E, and G to J). All data are mean ± SEM.

### DIANA response as non-BOLD effect

Despite convincing evidence presented here for DIANA’s ability to directly detect neuronal activity, it is possible that DIANA response might involve hemodynamic responses such as the BOLD effect. To dissect the BOLD effect from the DIANA response, BOLD-fMRI experiments were performed under two conditions: one condition as a default with extra oxygen-to-air ratio of 1:4 (oxygen:air condition) and the other condition with air only (Fig. 4A). The BOLD responses in the thalamus and S1BF following electrical whisker-pad stimulation in the air-only condition were significantly reduced compared to the oxygen:air condition (thalamus: 0.589 ± 0.045% to 0.251 ± 0.081%; contralateral S1BF: 1.128 ± 0.099% to 0.786 ± 0.127%, Fig. 4, B and C), consistent with BOLD activation’s dependence on oxygen supply (*33, 34*). In contrast, there was little change in the DIANA responses between air-only and oxygen:air conditions (thalamus: 0.188 ± 0.013 to 0.189 ± 0.009%; contralateral S1BF: 0.165 ± 0.009% to 0.168 ± 0.011%, Fig. 4, D and E). These results show that the hemodynamic BOLD response is not involved in the DIANA contrast mechanism.

**Fig. 4.**
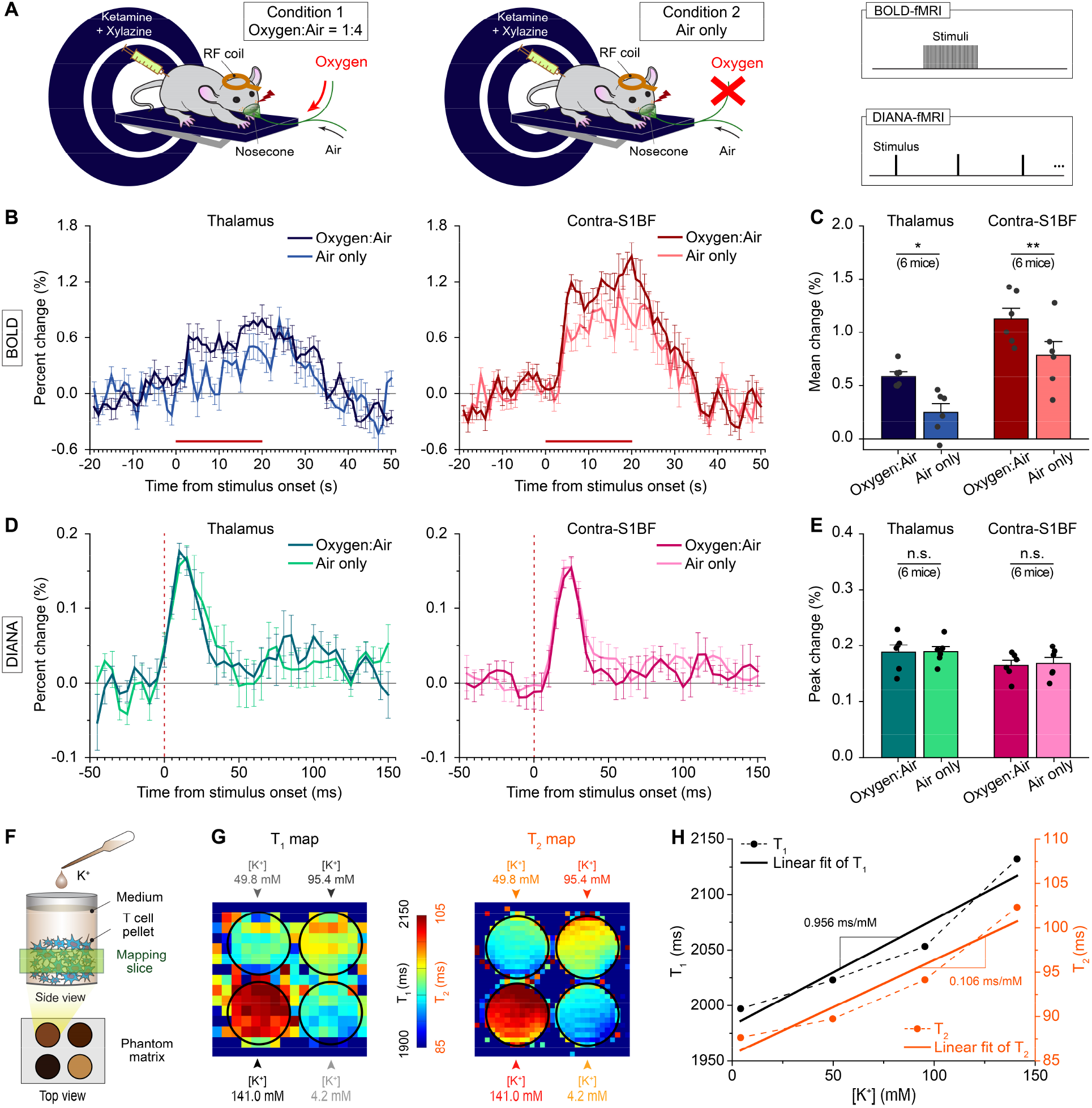
BOLD-independent DIANA response and its hypothesized contrast mechanism. (**A**) Illustration of oxygen challenge BOLD- and DIANA-fMRI experiments to image contralateral S1BF and thalamus applying electrical stimulation to left whisker pad in an anesthetized mouse on a 9.4 T scanner under two conditions: One condition as a default with a mixture of extra oxygen and air (1:4) (left) and the other condition with air only (middle) at the same flow rate. The same whisker-pad electrical stimulus (strength, 0.5 mA; pulse duration, 0.5 ms) was used for both BOLD- and DIANA-fMRI stimulation paradigms (right). (**B** and **C**) Percent signal changes of BOLD responses obtained from the thalamus (B, left) and the contralateral S1BF (B, right) under two conditions, and the corresponding bar graph showing the mean signal changes of BOLD responses in the thalamus and contralateral S1BF (C, *n* = 6 mice). The horizontal red bar indicates the period of electrical stimulation in BOLD-fMRI. (**D** and **E**) Same as (B and C) but with responses acquired using DIANA-fMRI (*n* = 6 mice). Vertical dotted lines indicate the stimulation onset time. (**F**) Illustration of a phantom experiment using T cells. (**G**) T_1_ and T_2_ relaxation times maps acquired at different extracellular K^+^ concentrations ([K^+^]) ranging from 4.2 to 141.0 mM. (**H**) Plots of T_1_ (black dotted line) and T_2_ (orange dotted line) changes with respect to [K^+^]. Solid lines are the linear fits to the T_1_ and T_2_ values to calculate the correlation coefficients with respect to [K^+^]. All data are mean ± SEM. *: *p* < 0.05, **: *p* < 0.01, n.s.: *p* > 0.05 for paired *t*-test.

Then what could be the possible signal source of DIANA response? Electromagnetic effects based on popular neuronal current models cannot be a candidate since signal loss caused by the phase cancellation of proton spins under the neuronal current-induced magnetic field change has a negative value (*3, 4, 9, 15*) while DIANA response has a positive signal change in the main lobe.

Instead, because changes in neuronal membrane potential can induce reorientation of the membrane interfacial water (*35*), DIANA response may arise from the changes in MR relaxation times (T_1_, T_2_) during neuronal activity, which is closely related to the amount of water molecules in the hydration layer on the membrane surface (*36*). To test our hypothesis, we performed phantom experiments using T cells to measure T_1_ and T_2_ relaxation times while T cells membrane potentials were manipulated by different extracellular K^+^ concentrations ([K^+^]) (*37*) (4.2 to 141.0 mM, Fig. 4F). We used T cells as phantom since, unlike neurons, T cells are homogeneous and immortal that can survive without oxygen supply *in vitro* in the scanner. T_1_ and T_2_ relaxation time maps (Fig. 4G) and values showed a strong positive correlation with [K^+^] (T_1_ slope, 0.956 ms/mM, R^2^ = 0.884; T_2_ slope, 0.106 ms/mM, R^2^ = 0.889, Fig. 4H). Thus, it is possible that the release of hydrated water into the free water around the cell membrane during membrane depolarization may account for T_1_ and T_2_ changes. Based on these measurements of T_1_ and T_2_ changes, Bloch simulation was performed to estimate DIANA signal changes and showed a positive signal change of ~0.13% in the main lobe (fig. S13), which is in good agreement with the experimental results (Figs. 1 to 3) and supports our hypothesis about the DIANA contrast mechanism.

## Discussion

Together, our results demonstrate that DIANA-fMRI indeed enables direct mapping of spike activity *in vivo* with high temporal (5 ms) and spatial (0.22 mm) resolution, as confirmed through *in vivo* electrophysiology combined with optogenetics. Such high temporal resolution of DIANA-fMRI even allowed the detection of the sequential propagation of neuronal activity through functionally-defined neural networks in the thalamocortical as well as cortico-thalamic pathways.

However, DIANA-fMRI has some limitations at this stage. Because its milliseconds temporal resolution is achieved based on line-scan data acquisition, DIANA-fMRI data is only generated in event-related responses to repetitive stimulation applications and, thus, it is challenging to investigate brain function in the resting state, or with a single stimulus. Moreover, DIANA-fMRI assumes that event-related responses in the brain are consistent across all the responses without trial-by-trial variability, which can somewhat undermine the reliability of DIANA-fMRI even with a modest change in neural spiking with each stimulation (*38*). Further investigation is also needed to fully elucidate the contrast mechanism of DIANA-fMRI.

There are interesting topics that can immediately be explored with DIANA-fMRI. One is to investigate whether DIANA-fMRI can noninvasively reveal rapid neural network dynamics in functionally-connected multiple, distant brain regions with a time scale of neural spike (*3, 39*). Furthermore, while we presented the DIANA-fMRI data detecting only fast-phasic neuronal activity here, by adjusting the inter-stimulation period long enough to measure the neuronal activity of interest, DIANA-fMRI could be used to measure diverse temporal patterns of neuronal activity on various time scales such as sustained tonic neuronal activity over a longer period of time. Another is to test its feasibility for human fMRI and translate DIANA-fMRI into a clinical human system. Approximate predictions only taking into account neuronal density, magnetic field strength, and typical voxel size in the animal and human systems suggest that DIANA-fMRI is likely to work in human studies as well (Supplementary Text). More complicated feedforward and feedback responses are expected to be observed in human brain networks (*40*), which could make DIANA response more challenging but even more interesting.

Overall, we expect DIANA-fMRI to open up new avenues in neuroimaging for a more accurate and deeper understanding of the brain’s functional organization, especially elucidating the causal relationship between temporal dynamics of neural networks and their function through the convergence of high temporal and spatial resolution.

## Acknowledgments

We thank Prof. Seong-Gi Kim’s group and Dr. Geunho Im for helping setting up mouse BOLD-fMRI experiments at the beginning of this study. We also thank Drs. Sungkwon Chung, Joonyeol Lee, and Choong-Wan Woo for valuable scientific discussion; Drs. Jae-Kyu Ryu and SoHyun Han for pulse sequence support; Chanhee Lee for MRI technical assistance; Seokwon Lee for administrative assistance.

## Funding

P.T.T., S.K.L, and J.Y.P. acknowledge financial support by the Brain Research Program through the National Research Foundation of Korea funded by the Ministry of Science, ICT, and Future Planning, Project ID NRF-2019M3C7A1031993. K.M. and J.L. acknowledge financial support from the same Brain Research Program, Project ID NRF-2019M3C7A1031994. H.J.J. and J.K acknowledge financial support from the same Brain Research Program, Project ID NRF-2019M3E5D2A01058328. P.T.T., S.K.L, and J.Y.P. also acknowledge financial support from the Institute for Basic Science (IBS-R015-D1).

## Author contributions

P.T.T. and J.Y.P. established the methodology for direct imaging of neuronal activity. P.T.T. prepared the pulse sequence, designed and conducted all animal MRI experiments. H.J.J. and J.K. designed all optogenetic experiments for MRI and electrophysiology. H.J.J. conducted the electrophysiological and optogenetic recordings, performed fluorescence and confocal imaging. H.J.J and J.K. analyzed electrophysiological data. K.M., S.K.L. and J.L. carried out MRI measurements of relaxation time changes in T cells and simulation. P.T.T., H.J.J., K.M., S.P.K., S.K.L., J.L., J.K., and J.Y.P. contributed to data analysis and discussion. P.T.T., H.J.J., J.K., and K.M. prepared the figures and drafts. P.T.T. and J.Y.P. wrote the paper. J.Y.P. conceived and supervised the project. All authors revised the manuscript.

## Competing interests

The authors declare no competing interests.

## Data and materials availability

The data can be provided upon reasonable request to corresponding authors. (J.Y.P.) and (J.K.).

## Supplementary Materials

Materials and Methods

Supplementary Text

Figs. S1 to S13

References (*41–52*)

## Notes

### Competing Interest Statement

The authors have declared no competing interest.

