## Supplementary Materials for "*In vivo* direct imaging of neuronal activity at high temporo-spatial resolution"

### Materials and Methods

Animals

C57BL/6 wild-type mice (23 – 33 g, 9 – 15 weeks old, Orient Bio and Daehan Biolink, South Korea) were used for both MRI and *in vivo* electrophysiology experiments with the approval of the Institutional Animal Care and Use Committee at Sungkyunkwan University (SKKUIACUC-2020-03-41-1) and Korea University (KUIACUC-2019-0068). Mice were maintained in a temperature-controlled environment on a 12h/12h light/dark cycle. Food and water were provided *ad libitum*.

MRI experiments

All MRI experiments were performed at 9.4 T (Bruker-BioSpec, 94/30 US/R) using an 86 mm inner-diameter volume coil for RF transmission and a 10 mm surface coil for signal reception.

*Animal preparation.* Mice were initially anesthetized under 4% isoflurane, then intraperitoneally (IP) injected with a mixture of 100 mg/kg ketamine and 10 mg/kg xylazine. Based on changes in physiological parameters during the experiment, supplementary doses (25 mg/kg ketamine and 1.25 mg/kg xylazine) were intermittently delivered to the mice (*41*). For all experiments except for the air-only condition in Fig. 4A, 1 liter/min flow of the oxygen:air (1:4) gases was continuously supplied to mice through a nosecone during the experiments, by which mice breathed spontaneously with two earplugs fixed and a bite bar on a customized cradle. Body temperature was maintained at 37 ± 0.5 °C by water heating system during the experiments. Motion-sensitive respiration (150 – 200 bpm) and electrocardiogram (ECG, 200 – 300 bpm) were continuously monitored with a physiological monitoring system (1030, Small Animal Instruments).

*Reference imaging.* As reference images for brain structures, twenty anatomical images of 0.5 mm coronal slices were acquired using two-dimensional (2D) fast low-angle shot (FLASH) imaging with scan parameters as follows: repetition time/echo time (TR/TE), 250/3 ms; flip angle (FA), 25°; field of view (FOV), 16 × 12 $mm^2$; image matrix, 256 × 192.

*BOLD-fMRI.* 2D gradient-echo echo-planar imaging (EPI) was conducted with scan parameters as follows: TR/TE, 1000/18 ms; FA, 60°; FOV, 16 × 12 $mm^2$; image matrix, 96 × 72; and slice thickness, 1 mm. Three adjacent coronal slices were acquired. After trimming the mouse whiskers bilaterally, electrical stimulation was delivered to the left whisker pad of the mice with 0.5 mA current strength, 0.5 ms pulse duration, and 4 Hz frequency using a five-electrode array connected to the isolator (ISO-Flex, A.M.P.I.) and pulse stimulator (Master 9, A.M.P.I). The stimulation paradigm consisted of 20 s pre-stimulation, 20 s stimulation, and 30 s post-stimulation.

*DIANA-fMRI.* 2D line-scan-based FLASH imaging was conducted to acquire the 5 ms temporal-resolution time series of 40 images in a single coronal slice, containing both thalamus and barrel field of primary somatosensory cortex (S1BF) (center at Anterior-Posterior (AP) -1.755 mm from bregma). Each TR consisted of a single line in the *k*-space during each interstimulus period, including slice-selective excitation with RF spoiling as well as phase- and frequency encodings. Scan parameters were as follows: TR/TE, 5/2 ms; FA, 4°; FOV, 16 × 12 $mm^2$; image matrix, 72 × 54; slice thickness, 1 mm; and scan time, 10.8 s/trial. The electrical stimulus of 0.5 ms duration was applied repeatedly at 200 ms interstimulus intervals (= 40 × 5 ms) which comprised 50 ms pre-stimulation and 150 ms post-stimulation including 0.5 ms stimulation duration, with 0.5 or 1 mA current strength.

In the postmortem experiment, the postmortem mice were acquired an hour after heart arrest induced by injecting saturated KCl bolus (50 – 80 µl) into the tail vein (Fig. 1, D and E).

In the sham experiment, an agar phantom was used with the same electrical stimulation scheme (0.5 mA) as in the *in vivo* mice experiment. Time series were extracted from three ROIs of a coronal slice (fig. S3).

Electrophysiological recording: electrical stimulation

Silicon probes were implanted in the contralateral S1BF and the thalamus to measure the neuronal activities in response to the same electrical stimulation of the mouse whisker-pad as in DIANA-fMRI. The mouse head was fixed to a stereotaxic device (51730, Stoelting Inc.) under anesthesia (ketamine, 75-100 mg/kg, and medetomidine, 1 mg/kg). After trimming the mouse whiskers bilaterally, the left whisker-pad of the mouse was electrically stimulated using a five-electrode array which delivered a hundred current pulses of 0.5 ms duration and 0.5 mA amplitude repeatedly at 200 ms interstimulus intervals. Electrical stimulation was given using an isolated current stimulator (DS3, Digitimer) controlled by a BioAmp Processor (RZ2 system, Tucker-Davis Technology). A 32-channel silicon probe (A1×32-poly2-5mm-50s-177-OA32, Neuronexus) was implanted into the right S1BF (AP -1.94 mm, Medial-Lateral (ML) +3.3 mm from bregma; 850-900 µm depth from pia, at an angle of 25-30° from the vertical to a probe tip) and, when needed, an additional 16-channel silicon probe (A1x16-10mm-100-177-A16, Neuronexus) was implanted into the right thalamus (AP -1.94 mm, ML +1.5 mm from bregma; 3.3 mm depth from pia). Electrophysiological signals were sampled at 25 kHz and used for analyzing the single unit activities and LFP thereof. To determine the position of each probe, silicon probes were coated with a 1,1′-Dioctadecyl-3,3,3′,3′-Tetramethylindocarbocyanine Perchlorate (DiI) fluorescent dye before insertion of the probes. Body temperature was monitored and maintained at 37 °C using a DC temperature control system (40-90-8D, FHC Inc.) in all experiments.

Virus and stereotaxic surgery for optogenetics

A blue light-gated cation channel (Channelrhodopsin-2, ChR2) (*42*) was expressed in the excitatory neurons of S1BF by injecting adeno-associated virus (rAAV) vectors AAV5-CaMKII-hChR2(E123T/T159C)-p2A-mCherry-WPRE (UNC Vector Core) in solution (150-300 nl, $3.8\times10^{12}$ virus molecules/ml) to the right S1BF (AP -1.94 mm, ML +3.3 mm from bregma) while the mouse head was fixed to a stereotaxic device (51730, Stoelting Inc.) under isoflurane anesthesia. Virus injections were made at two cortical depths (300 and 600 µm depth from pia) using a 5 µl syringe with a blunt tip connected to a motorized stereotaxic injector (Stoelting Quintessential Injector, 53311, Stoelting Inc.) at a speed of 30-50 nl/min. To prevent the withdrawal of the virus and allow for virus diffusion, the injecting needle remained in the brain for more than five minutes after the injection. For *in vivo* optogenetic stimulation of ChR2-expressing excitatory neurons in the S1BF during the MRI scan, a fiber-optic cannula (200 µm diameter, 0.22 NA, Thorlabs) was placed at an angle of 25 – 30° to the pia of S1BF (AP -1.94 mm, ML +3.3 mm from bregma) and was secured to the skull using resin dental cement (Vertex Self-

Curing, Vertex Dental). A minimum of two weeks recovery time was required before performing optogenetic stimulation *in vivo*.

Optogenetic DIANA-fMRI

Cortical excitatory neurons were activated in the S1BF by expressing ChR2 and stimulating them with blue light (473 nm). A time series of 50 images were acquired at the same slice position as in DIANA-fMRI mentioned above. Neuronal activities were investigated not only in the S1BF but also in the thalamus. A fiber-optic cannula implanted chronically in the mouse head was connected to a blue light (473 nm) laser (BL473T8U-150FC, SLOC) outside the scanner room via an optical fiber. To see the dependence of the neuronal responses under varying stimulation conditions, two different durations (20 and 50 ms) at 50 $mW/mm^2$ blue light intensity were used with 250 ms interstimulus period (= 50 × 5 ms). The interstimulus period consisted of 50 ms pre-stimulation, 20 or 50 ms stimulation, and 180 or 150 ms post-stimulation.

Electrophysiological recording: optogenetic stimulation

The same optogenetic stimulation scheme as in the optogenetic DIANA-fMRI was used for electrophysiological recording *in vivo*. A blue light (473 nm) laser (iBeam-smart-473, Toptica Photonics) was used to activate ChR2-expressing excitatory neurons in the S1BF and the electrophysiological recordings were performed in both S1BF and thalamus simultaneously. The blue light was transmitted through an optical fiber (200 µm diameter, 0.22 NA) stacked on a 32-channel silicon probe (A1x32-poly2-5mm-50s-177-OA32, Neuronexus). The same blue light stimulation as in the optogenetic DIANA-fMRI was applied, but with 50 repetitions of blue light stimulation.

Fluorescence imaging

To confirm the location of the silicone probe and the expression of ChR2 opsins in excitatory neurons of S1BF, mice were decapitated under isoflurane anesthesia and the brains were transferred to a vibratome chamber which was filled with ice-cold artificial cerebrospinal fluid containing 126 mM NaCl, 3 mM KCl, 1.25 mM $NaH_2PO_4$, 2 mM $MgSO_4$, 2 mM $CaCl_2$, 25 mM $NaHCO_3$, and 10 mM glucose at pH 7.2 – 7.4. Coronal brain slices (300 µm) were cut using a vibratome (VT1000s, Leica) and fixed overnight in 4% paraformaldehyde (PFA, Sigma-Aldrich) for longer than 24 hours at 4°C. Fixed slices were washed three times in wash buffer (0.3% Triton X-100 in 0.1M PBS) and mounted on glass slides with mounting medium (H-1200-NB, Vector Lab). Fluorescent signals and images of expressing ChR2 opsins were acquired using confocal microscopy (LSM-700, Zeiss) and fluorescent images of the silicone probes were obtained using fluorescent microscopy (DM-2500, Leica).

Measurement of changes in $T_1$ and $T_2$ relaxation times

An immortalized line of human T-lymphocyte cells (T cells) was used as an experimental model *in vitro*. They were cultured in Roswell Park Memorial Institute (RPMI) 1640 medium with 10% fetal bovine serum and 1% penicillin/streptomycin (Merck/Sigma Aldrich). The cultured cells were evenly distributed in four cylindrical cavities machined into a square acrylic phantom and centrifuged to a pellet form immediately before MRI acquisition. To manipulate the resting membrane potential by depolarization, four different concentrations of potassium ions ([$K^+$], 4.2, 49.8, 95.4, and 141.0 mM) were applied to each of the cavities that contain T cell pellet (*37*). For $T_1$ estimation, the inversion recovery sequence with EPI readout was used (image matrix, 32 × 16). $T_1$ values were obtained voxelwise by fitting magnitude data to an exponential recovery function. For $T_2$ estimation, a multi-echo spin-echo sequence was used (image matrix, 64 × 32) and $T_2$ values were obtained voxelwise by fitting magnitude data to an exponential decay function. The region of interest (ROI) was set as a circle in the center of each cavity containing the T cell pellet.

Simulation of DIANA signal change

Based on the measured $T_1$ and $T_2$ values, the MR signal change was estimated by Bloch simulations (*43*) using the same pulse-sequence scheme as well as the same scan parameters (TR, 5 ms; TE, 2 ms; FA, 4°) as in DIANA-fMRI mentioned above. A perfect spoiling of the transverse magnetization was assumed immediately before each RF pulse, and signals were acquired after the spin system reached a steady state. A time series of 40 single free induction decays (FIDs) were emulated with the transverse dephasing time set to $T_2$ (assumed to be the same as $T_2^*$). Neuronal firings were assumed to be evoked not simultaneously but by a Gaussian distribution (standard deviation, 0.78 ms) so that the average membrane potential of the neurons at its most depolarized moment was 0 mV. The shape of membrane potential over time was assumed to follow that of the action potential and simulated by using the Hodgkin-Huxley model with the following parameters: $K^+$ Nernst potential ($V_k$), -77 mV; $Na^+$ Nernst potential ($V_{Na}$), 50 mV; resting potential ($V_r$), -65 mV; membrane capacitance per unit area ($c_m$), 0.01 F/m$^2$; axon radius ($a$), 5 μm; membrane thickness ($b$), 6 nm; axoplasm resistivity ($\rho_i$), 1.1 Ω·m. With respect to [$K^+$], 4.2 mM and 141.0 mM were assumed to correspond to the resting state and the most depolarized state, respectively. Accordingly, for the resting state, $T_1$ and $T_2$ values were set as 1996.99 and 87.64 ms, respectively, and, for the most depolarized state, they were set as 2132.23 and 102.33 ms, respectively (Fig. 4H). During the period of neural spiking activity, the $T_1$ and $T_2$ values were assumed to change linearly with the action potential.

Data analysis of mouse fMRI

For all analyses of DIANA- and BOLD-fMRI data, brain structures, and ROI definitions were referenced in the Allen Mouse Brain Atlas (*44*) (Allen Brain Institute, http://mouse.brain-map.org). All the data was processed using home-built MATLAB codes (R2019a/b, MathWorks) with additional help of some external programs for BOLD-fMRI data such as Analysis of Functional Neuroimages package (AFNI) (*45*), FMRIB Software Library (FSL) (*46*), and Advanced Normalization Tools (ANTs) (*47*).

*BOLD-fMRI.* Each mouse data was obtained from an average of 5 to 10 trials. For BOLD responses to the electrical stimulation ($n$ = 10), time courses were extracted from circular ROIs of 0.5 mm radius (including 29 voxels) in the thalamus and S1BF areas. In particular, the defined ROIs based on the atlas were applied to the oxygen challenge data. For comparison of the BOLD responses between different ventilation conditions, the average response was calculated by averaging the BOLD percent changes from $6^{th}$ to $20^{th}$ seconds after stimulation onset. Individual BOLD activation maps were co-registered and overlaid on the FLASH images with a statistical threshold of uncorrected $p < 0.05$ and cluster size > 5 voxels. To obtain a group-averaged BOLD map, individual EPI images were normalized to a coronal template which was made by averaging the individual FLASH images. The group-averaged BOLD maps were generated using one-sample *t*-test with uncorrected $p < 0.05$ criterion. All the maps were spatially smoothed using a Gaussian kernel of 0.2 mm full width at half maximum (FWHM).

*DIANA-fMRI.* Two signal processing steps were taken to obtain the DIANA time series. Firstly, temporal smoothing was applied using a 15 ms Gaussian kernel (three-point kernel) before extraction of DIANA signals from the defined ROIs in the S1BF (30 – 35 voxels) and thalamus (69 voxels). Linear detrending was then applied for the removal of signal drift. The percent signal change defined as $\Delta S/S_0*100$ was calculated to represent the DIANA signal change, where $\Delta S$ is a signal difference with reference to the baseline ($S_0$) that was given by averaging the signal of the pre-stimulation period, i.e., from $1^{st}$ to $10^{th}$ images in the time series. All the time series data was analyzed after the mean was taken across all the trials as well as animals: for DIANA-fMRI data using the electrical stimulation (including the control), 40 trials/mouse except for the postmortem data (30 trials/mouse); for DIANA-fMRI data using the optogenetic stimulation, 13 – 20 trials/mouse. To show the spatial distribution of DIANA responses, *t*-value maps were generated from the trials for individual mice. Prior to the activation mapping, an extra spatial smoothing was applied using a two-dimensional median filter with a square kernel of 5 voxels to highlight the responsive areas. During the activation mapping, each image in the post-stimulation period (i.e., $11^{th}$ to $40^{th}$ images in the time series with electrical stimulation, $11^{th}$ to $50^{th}$ images with optogenetic stimulation) was compared voxelwise with reference to the baseline (defined as an average of the pre-stimulation signals) using the paired *t*-test. Positive *t*-value voxels were displayed and overlaid on the original DIANA images, provided that the statistical criteria of $p < 0.05$ and cluster size > 5 voxels were met.

### Data analysis of electrophysiology

The single-unit spike activities were identified from the multi-unit activities (MUA) by applying a band-pass filter (0.3 to 5 kHz) to the recorded electrophysiological signals. To detect and sort out the spikes, a KiloSort software (*48*) was used with a threshold level of 4.5 times the standard deviation of each MUA. As a single unit, some units among the detected ones were adopted that clearly showed the negative deflection in the spike waveform and did not violate the refractory period (2 – 3 ms) in the auto-correlogram of spike times (*49*). Since only subsets of cortical neurons respond to sensory input (*50*) and DIANA-fMRI is assumed to measure signals from the neurons undergoing depolarization of the membrane potential, only the whisker-pad stimulation-responsive single units were used for the analysis. A single unit was considered stimulation-responsive provided that its spike firing rate increased during the 50 ms post-stimulation period when compared with the 100 ms pre-stimulation period, at a statistical threshold of $p < 0.05$ using the Mann-Whitney U-test. To investigate the correlation of single-unit activity with DIANA response, the latency of the peak spike firing rate was analyzed from the post-stimulus time histogram (PSTH). To see whether local field potential (LFP) also correlates with the DIANA signal, the recorded LFPs were filtered using a band-pass filter (0.1 to 100 Hz) and downsampled to 1 kHz. The delay and amplitude of LFP were measured as the difference between the negative peak of LFP during 100 ms after stimulation and the baseline calculated as the average of LFP during 10 ms before stimulation (*51*).

Statistics

All statistical comparisons between the two groups were conducted using two-tailed *t*-test (paired or unpaired *t*-test), and one-way analysis of variance (ANOVA) with Bonferroni *post hoc* test was used for comparing three groups. Quantitative data were all presented as mean ± standard error of the mean (S.E.M.). Normality was tested with D'Agostino's K-squared method except for the samples whose number exceeds 30.

### Supplementary Text

Although the field strengths of typical human MRI systems are lower than those of animal systems, e.g., ranging from 1.5 T to 7 T, successful translation into human fMRI is conceivable based on signal-sensitivity consideration. It is well known that there are about 86 billion neurons in the human brain, with an average weight of 1,500 g, and about 70 million neurons in the mouse brain, with an average weight of 0.4 g, which correspond to $5.73 \times 10^7$ neurons/g and $1.75 \times 10^8$ neurons/g in terms of neurons per unit mass, respectively (*52*). Considering that the signal-to-noise ratio (SNR) increases linearly with the field strength to a good approximation and that the number of neurons in the mouse brain per gram is about three times that in the human brain, DIANA-fMRI is expected to show about ten times greater signal sensitivity in animal studies at 9.4 T than human studies at 3 T when the brain tissue density is assumed to be the same. Fortunately, the voxel size of human brain images is typically much larger than that of mouse brain images, e.g., by two hundred times in the case of typical human BOLD-fMRI (e.g., $2\times2\times2$ $mm^3$) versus animal DIANA-fMRI (e.g., $0.2\times0.2\times1$ $mm^3$), so the signal sensitivity can be about twenty times greater in the human study, making up for the loss of signal sensitivity caused by lower field strengths, less neuronal density, and larger coil size. On the other hand, some negative effects such as motion artifact will be more serious in the human studies compared to the animal studies.

**Supplementary Figures**

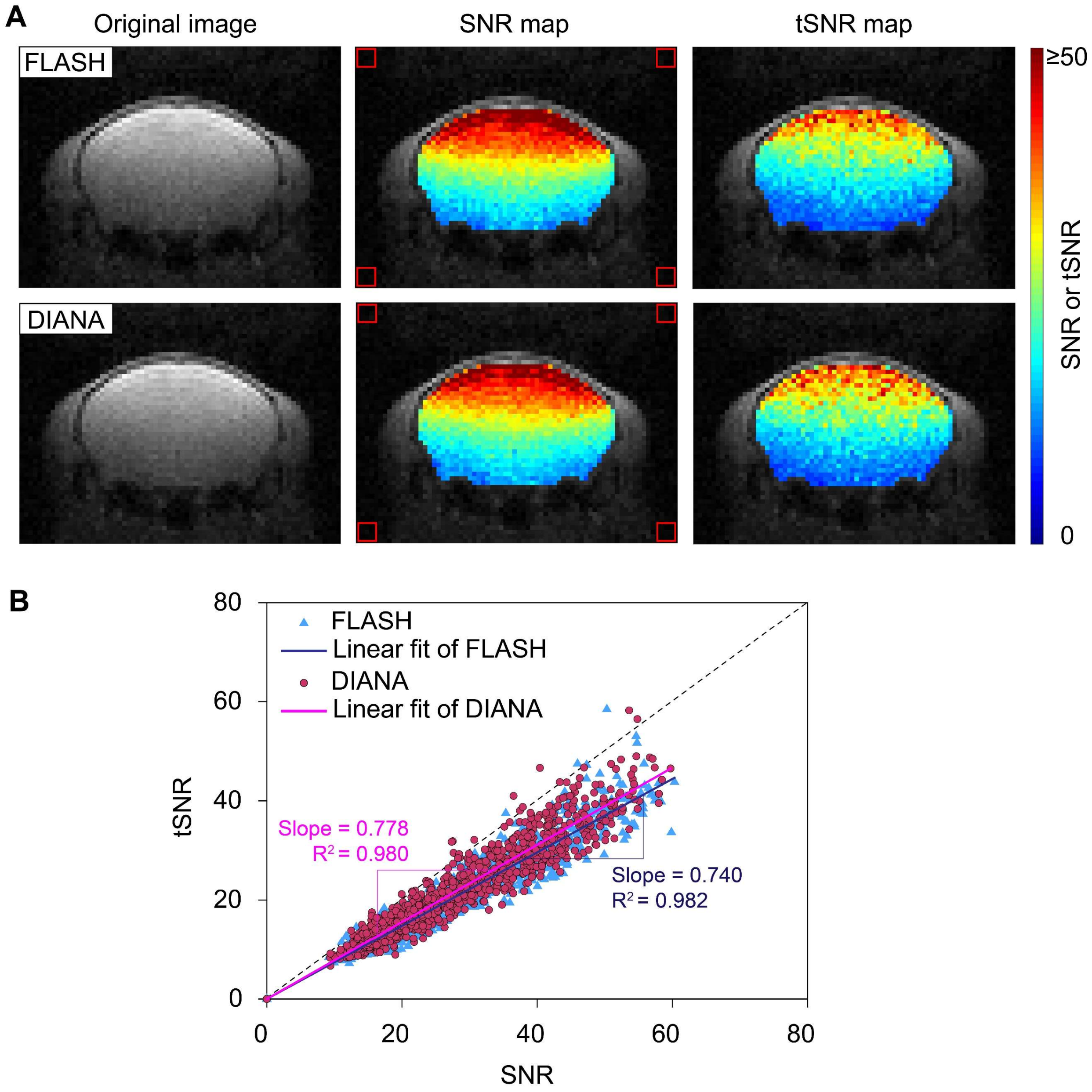


**Fig. S1. Comparison of SNR and tSNR between DIANA-fMRI and FLASH imaging.** (**A**) Voxelwise SNR and tSNR maps of FLASH imaging (top) and DIANA-fMRI (bottom) of an *in vivo* mouse brain in a single trial. The SNR map was generated from the first image in the time series by dividing each voxel by the standard deviation of background noises defined at four corners of the image (red square boxes). The tSNR map was calculated by dividing the voxelwise average of the time series by its temporal fluctuation. (**B**) A scatter plot of SNR versus tSNR values for FLASH imaging (blue triangular dots) and DIANA-fMRI (magenta circular dots). Solid lines indicate the linear fits of SNR/tSNR scatters for FLASH imaging (dark blue) and DIANA-fMRI (light magenta).

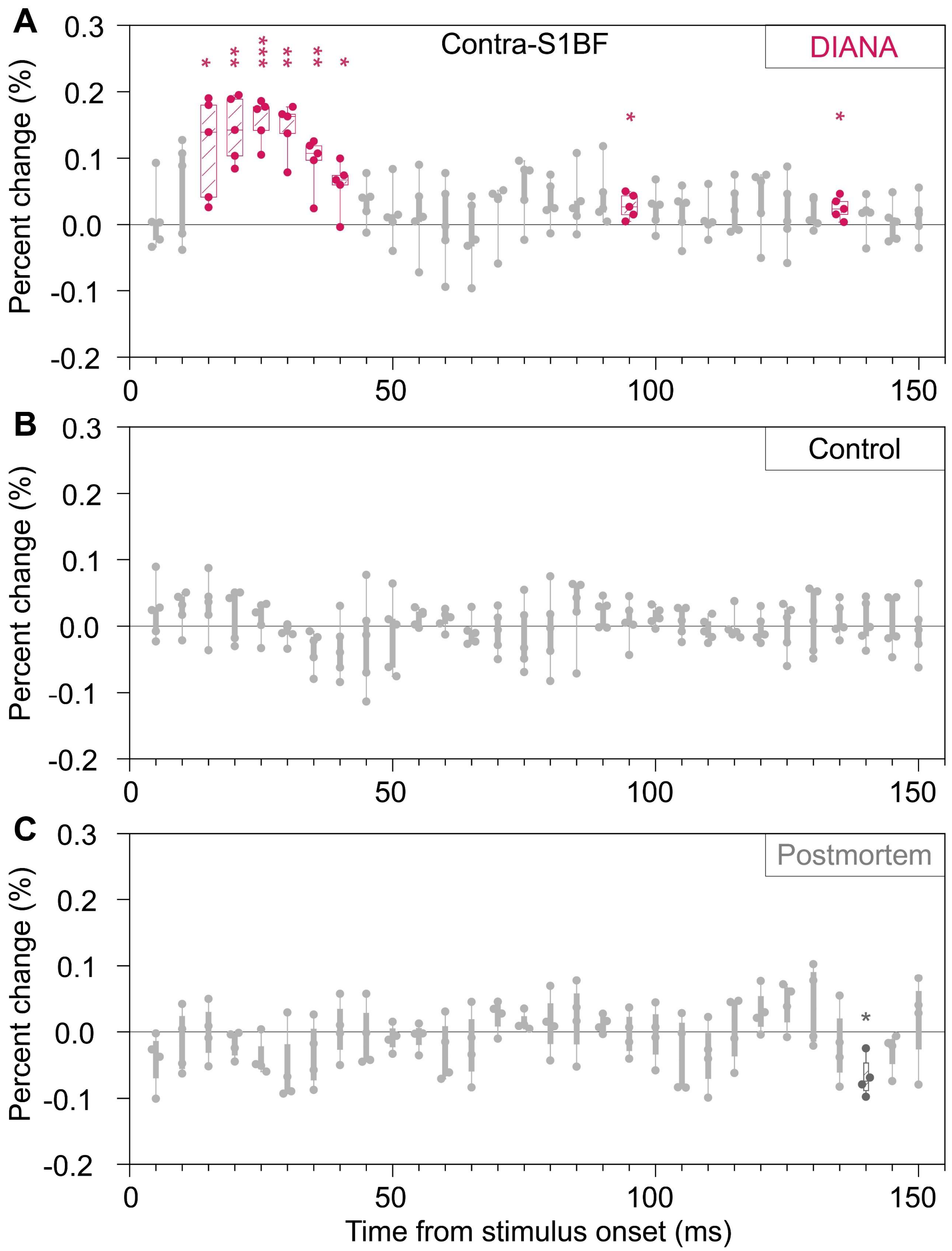


**Fig. S2. Time-dependent statistical significance of DIANA signal changes.** (**A** to **C**) DIANA responses in the right barrel field of primary somatosensory cortex (S1BF) applying electrical stimulation of left whisker pad in anesthetized mice on a 9.4 T scanner ($n$ = 5 mice) (A), without stimulation in the control mice ($n$ = 5 mice) (B), and with stimulation in the postmortem mice ($n$ = 4 mice) (C). In the box plots, each box represents 25$^{th}$ to 75$^{th}$ percentiles, horizontal lines represent the median, and whiskers range from the minimum to the maximum values. *: $p < 0.05$, **: $p < 0.01$, ***: $p < 0.001$ for paired $t$-test.

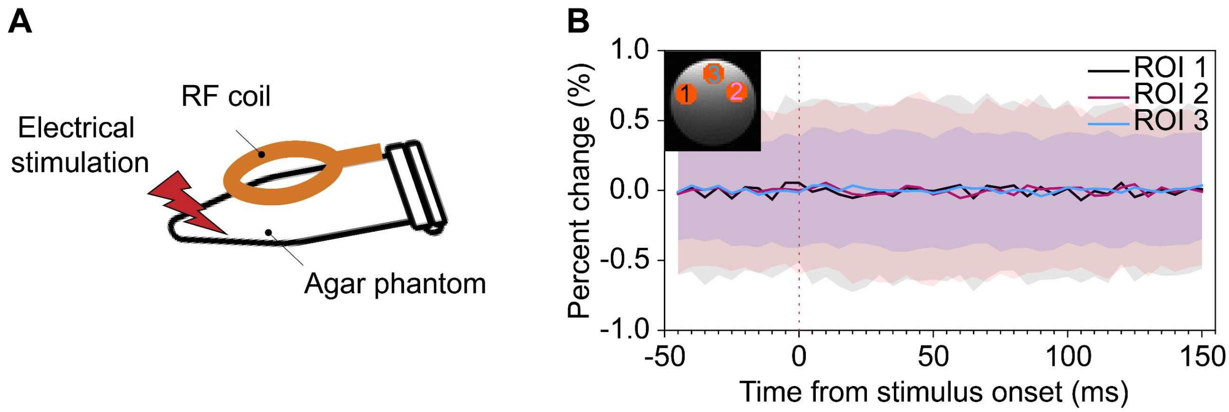


**Fig. S3. Sham DIANA-fMRI experiment.** (**A**) Illustration of a sham DIANA-fMRI experiment with electrical stimulation using an agar phantom instead of an *in vivo* mouse on a 9.4 T scanner. (**B**) Time series of percent signal changes in DIANA responses extracted from three regions-of-interest (ROI, defined in the subfigure) in a 1 mm coronal slice. Data from 400 trials were averaged, where the number of trials was determined by considering the group-averaged results of mice DIANA-fMRI, e.g., 40 trials/mouse × 10 mice in Fig. 2D, for comparison of temporal fluctuation. The red vertical dotted line indicates the electrical stimulation onset time. Mean values were displayed as solid lines and standard deviations in shades (black, ROI 1; magenta, ROI 2; blue, ROI 3).

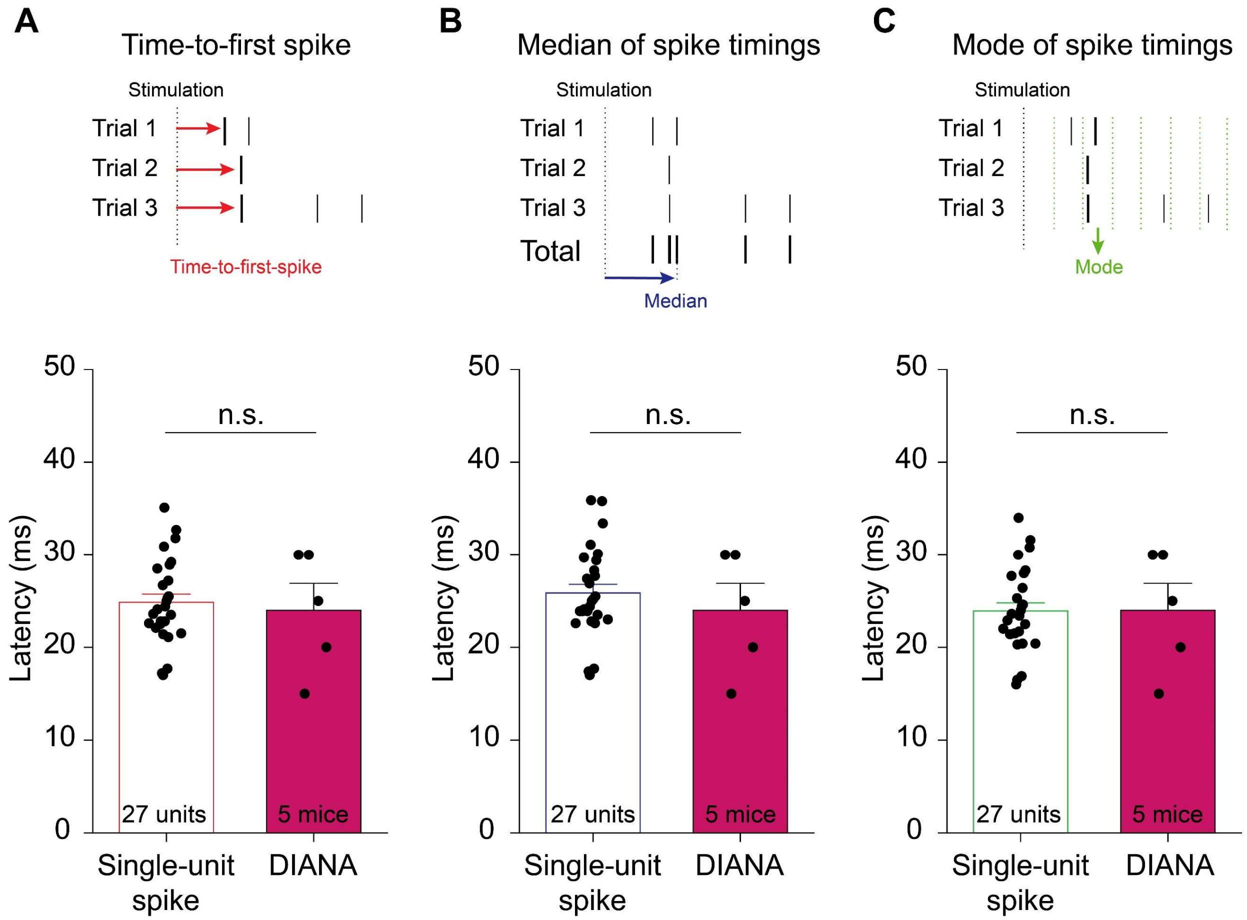


**Fig. S4. Other temporal spike characteristics of whisker-pad stimulation-responsive single units also match the DIANA response latency.** Top: Illustration of time-to-first-spike analysis (**A**), median of spike timings (**B**), and mode of spike timings (**C**). Bottom: Latency of time-to-first spikes (A), median (B) and mode (C) of whisker-pad stimulation-responsive spike timings in contralateral S1BF ($n$ = 27 units from 10 mice), in comparison to the latency of peak DIANA responses ($n$ = 5 mice). Vertical dotted lines indicate the stimulation onset time. All data are mean ± SEM. n.s.: $p > 0.05$ for unpaired $t$-test.

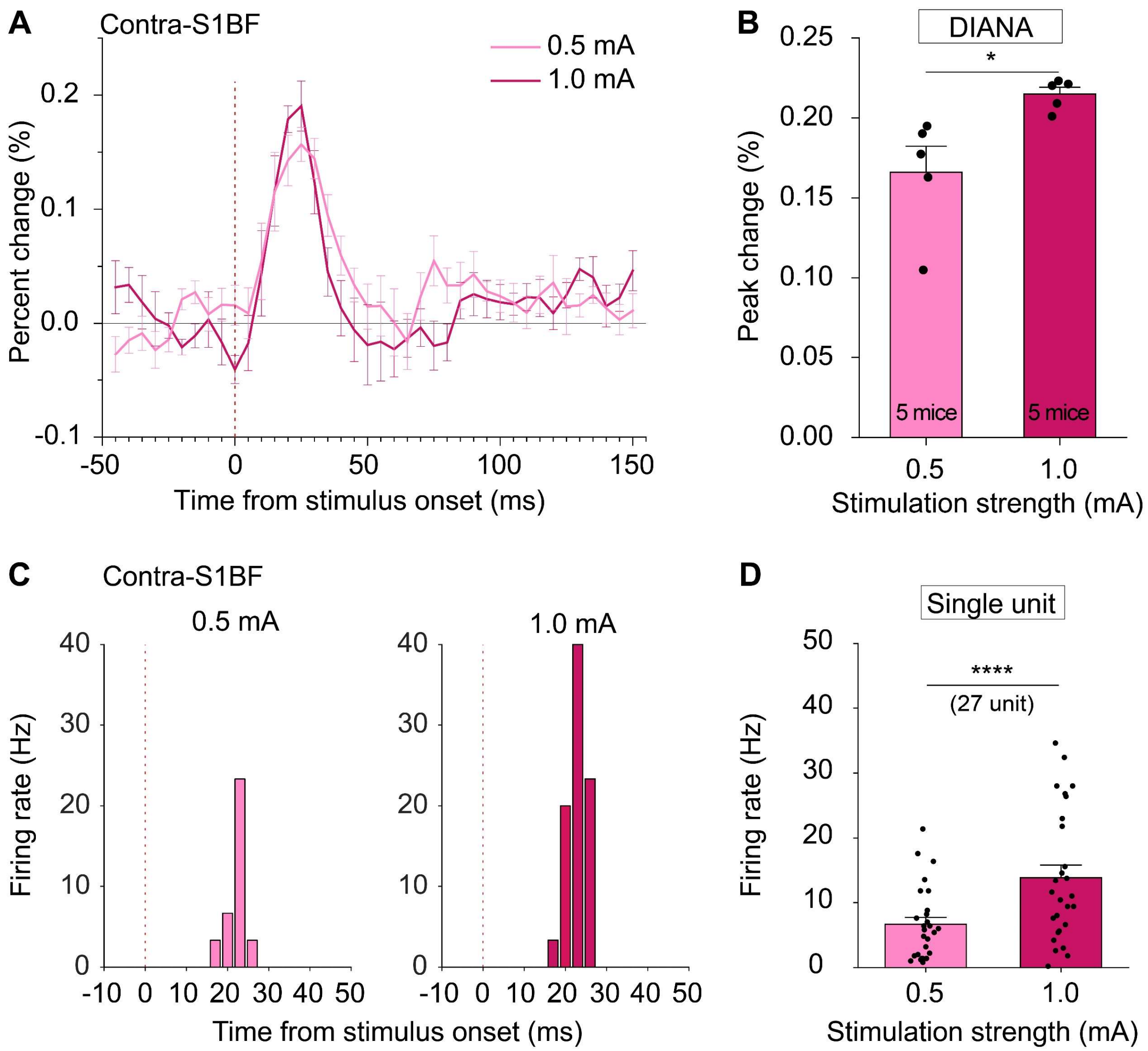


**Fig. S5. Dependence of DIANA responses and spike firing rates on the strength of electrical whisker-pad stimulation.** The time series (**A**) and percent changes (**B**) of peak DIANA responses in the contralateral S1BF in response to 0.5 mA (light magenta) and 1 mA electrical whisker-pad stimulation (magenta) ($n$ = 5 mice). Post-stimulation time histogram (PSTH) (**C**) of the whisker-pad stimulation-responsive single units over time for 0.5 mA (left) and 1 mA (right) current strength, and bar graph showing their spike firing rates (**D**) ($n$ = 27 units from 10 mice). Vertical dotted lines indicate the stimulation onset time. All data are mean ± SEM. *: $p < 0.05$, ****: $p < 0.0001$ for paired $t$-test.

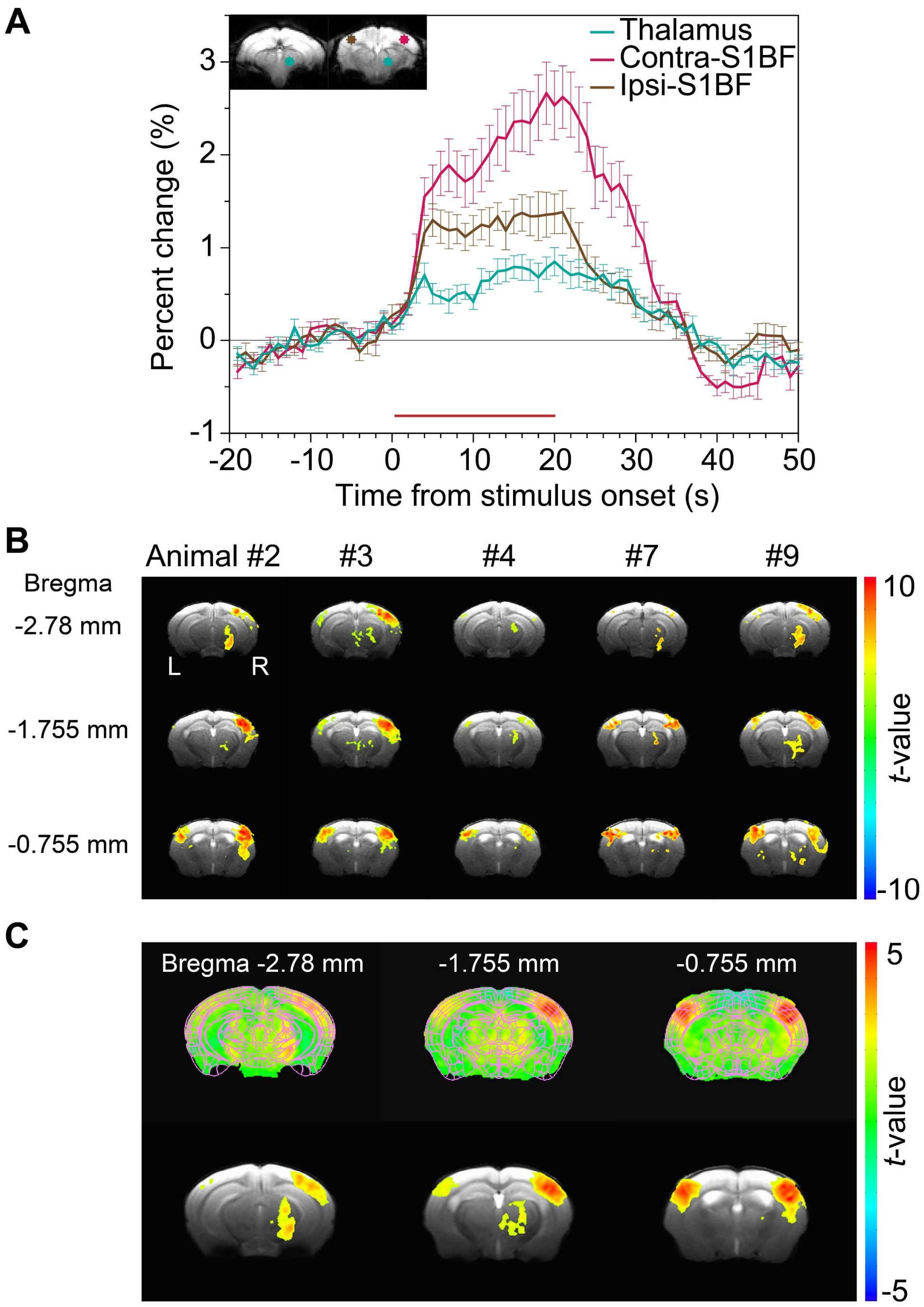


**Fig. S6. Blood-oxygenation-level-dependent (BOLD) activations in response to electrical whisker-pad stimulation.** (**A**) Percent signal changes of BOLD responses in the thalamus, contra- and ipsilateral S1BF (defined ROIs in subfigures, $n = 10$ mice). The horizontal red bar indicates the duration of electrical whisker-pad stimulation. (**B**) Individual BOLD activation maps of 5 representative mice (uncorrected $p < 0.05$, cluster size > 5 voxels). (**C**) Group-averaged activation maps without (top) and with (bottom) statistical thresholds (one-sample *t*-test, uncorrected $p < 0.05$). All data are mean ± SEM.

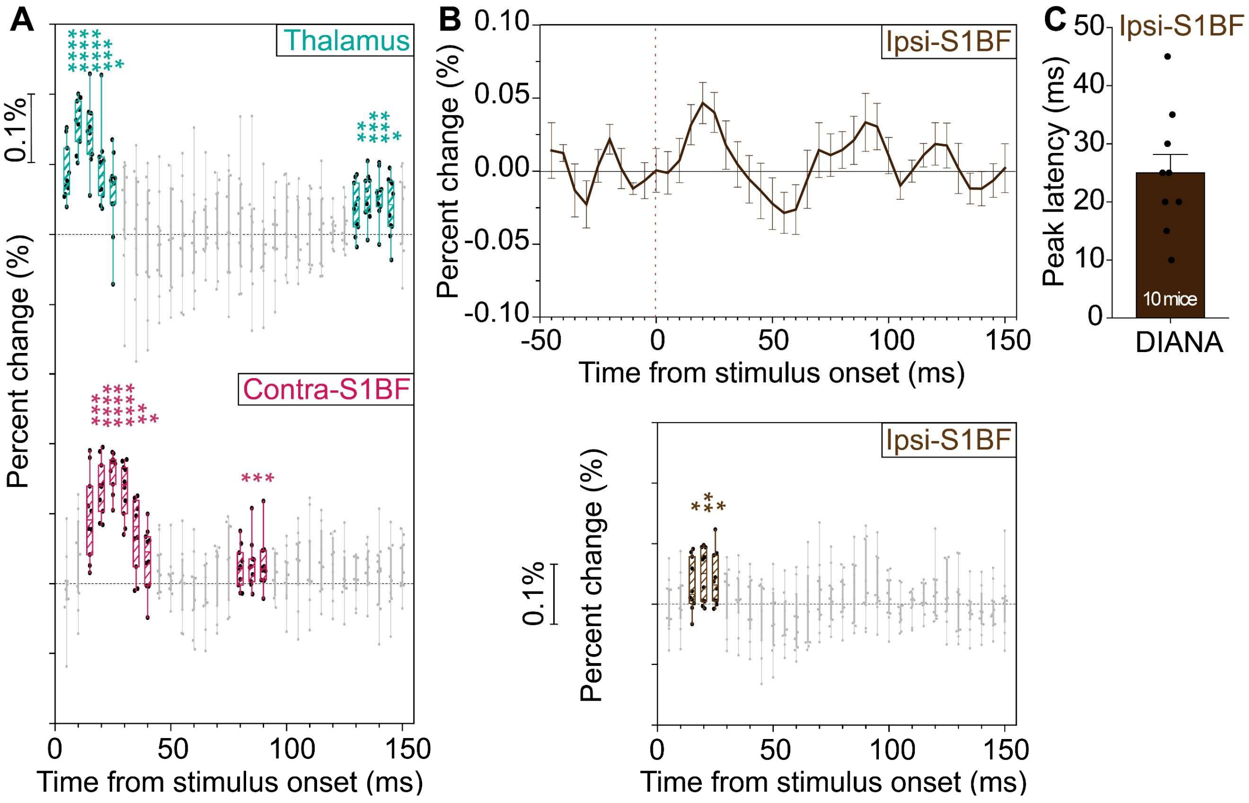


**Fig. S7. Time-dependent statistical significance of DIANA signal changes in the thalamus, contralateral and ipsilateral S1BF.** (**A**) Time-dependent statistical significance of DIANA responses in the thalamus (top) and contralateral S1BF (bottom) in response to electrical whisker-pad stimulation ($n$ = 10 mice). (**B**) Percent signal changes of DIANA responses (top) and their time-dependent statistical significance (bottom) in the ipsilateral S1BF. (**C**) Bar graph showing the latency of peak DIANA responses in the ipsilateral S1BF. The vertical dotted line in (B) indicates the whisker-pad stimulation onset time. In the box plots, each box represents 25$^{th}$ to 75$^{th}$ percentiles, horizontal lines represent the median, and whiskers range from the minimum to the maximum values. All data are mean ± SEM. *: $p < 0.05$, **: $p < 0.01$, ***: $p < 0.001$, ****: $p < 0.0001$ for paired $t$-test.

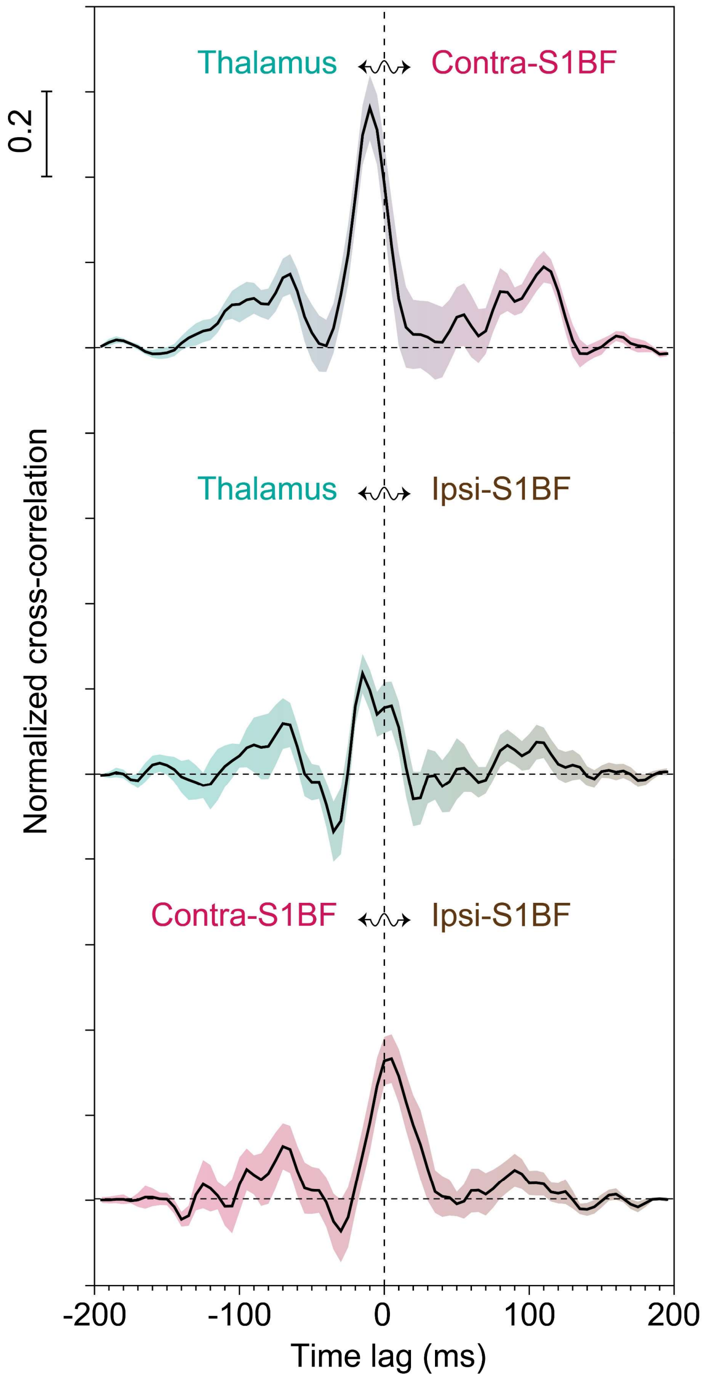


**Fig. S8. Further cross-correlation analysis of DIANA-fMRI signals.** Cross correlation of DIANA-fMRI signals from different areas responding to electrical whisker-pad stimulation: between thalamus and contralateral S1BF (top), thalamus and ipsilateral S1BF (middle), and contralateral and ipsilateral S1BF (bottom) ($n$ = 10 mice). All data are mean ± SEM.

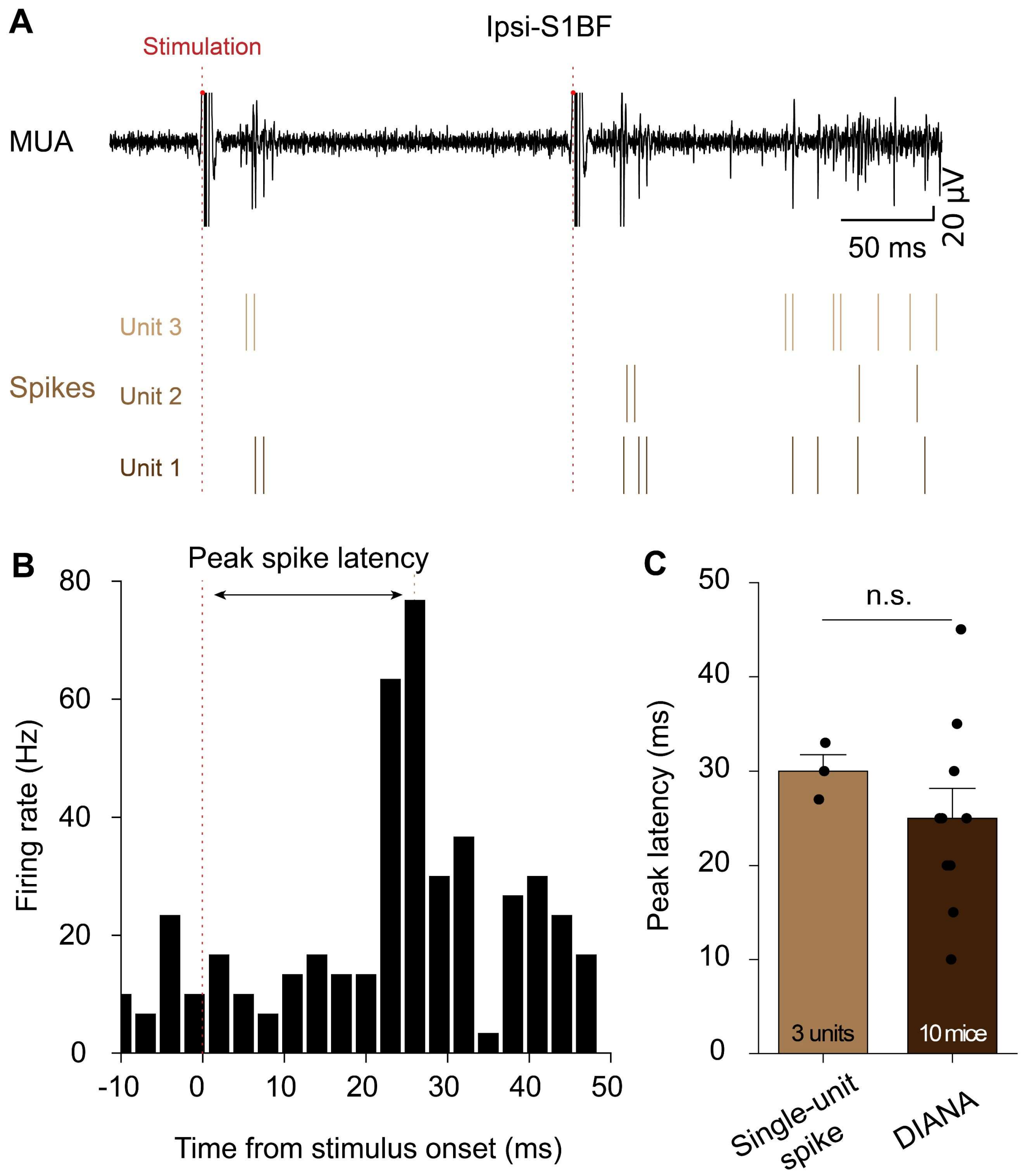


**Fig. S9. Electrophysiological recording in ipsilateral S1BF with electrical whisker-pad stimulation *in vivo*.** (**A**) A representative multi-unit activity (MUA) (black trace, top) from which single-unit spikes (brown, bottom) were analyzed in the ipsilateral S1BF. (**B**) Post-stimulation time histogram (PSTH) of the whisker-pad stimulation-responsive single units over time in the ipsilateral S1BF. (**C**) Bar graph showing the latencies of peak spike firing rates of the whisker-pad stimulation-responsive single units ($n$ = 3 units from 1 mouse) and DIANA responses ($n$ = 10 mice). Vertical dotted lines indicate the whisker-pad stimulation onset time (red, A and B) and latency of peak spike firing rates (brown, B). All data are mean ± SEM. n.s.: $p > 0.05$ for unpaired $t$-test.

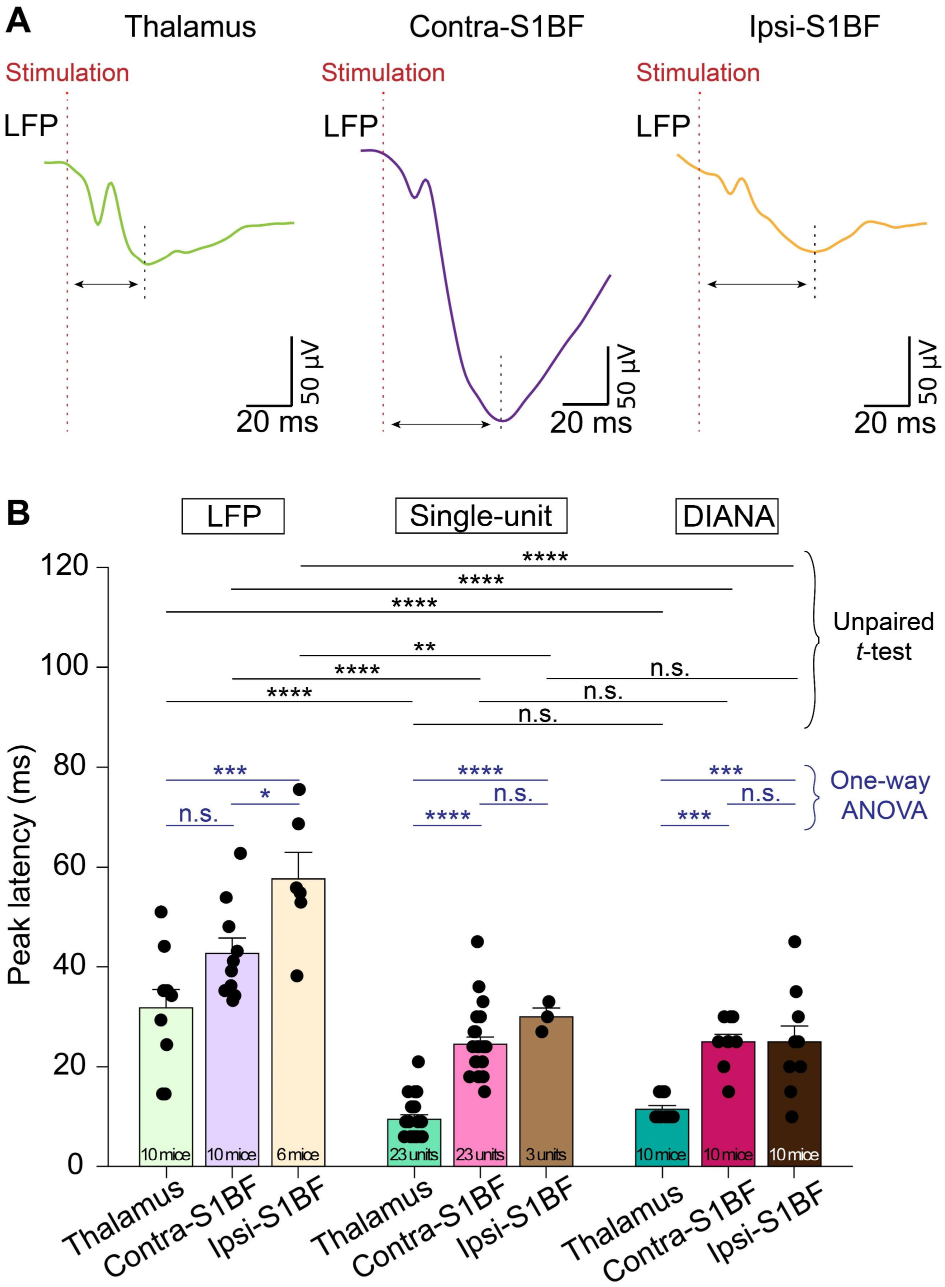


**Fig. S10. Comparison of DIANA response to local field potential (LFP) and single-unit spike in response to electrical whisker-pad stimulation *in vivo*.** (**A**) Representative LFPs in the thalamus (left), contralateral S1BF (middle), and ipsilateral S1BF (right). (**B**) Bar graph showing the latencies of peak LFPs (thalamus and contralateral S1BF, $n = 10$ mice; ipsilateral S1BF, $n = 6$ mice) compared to the latencies of peak spike firing rates (thalamus, $n = 23$ units from 10 mice; contralateral S1BF, $n = 23$ units from 5 mice; ipsilateral S1BF, $n = 3$ units from 1 mouse) and peak DIANA responses ($n = 10$ mice). Vertical dotted lines indicate the whisker-pad stimulation onset time (red) and LFP peaks (black). All data are mean ± SEM. *: $p < 0.05$, **: $p < 0.01$, ***: $p < 0.001$, ****: $p < 0.0001$, n.s.: $p > 0.05$ for one-way ANOVA with Bonferroni *post hoc* test and unpaired *t*-test.

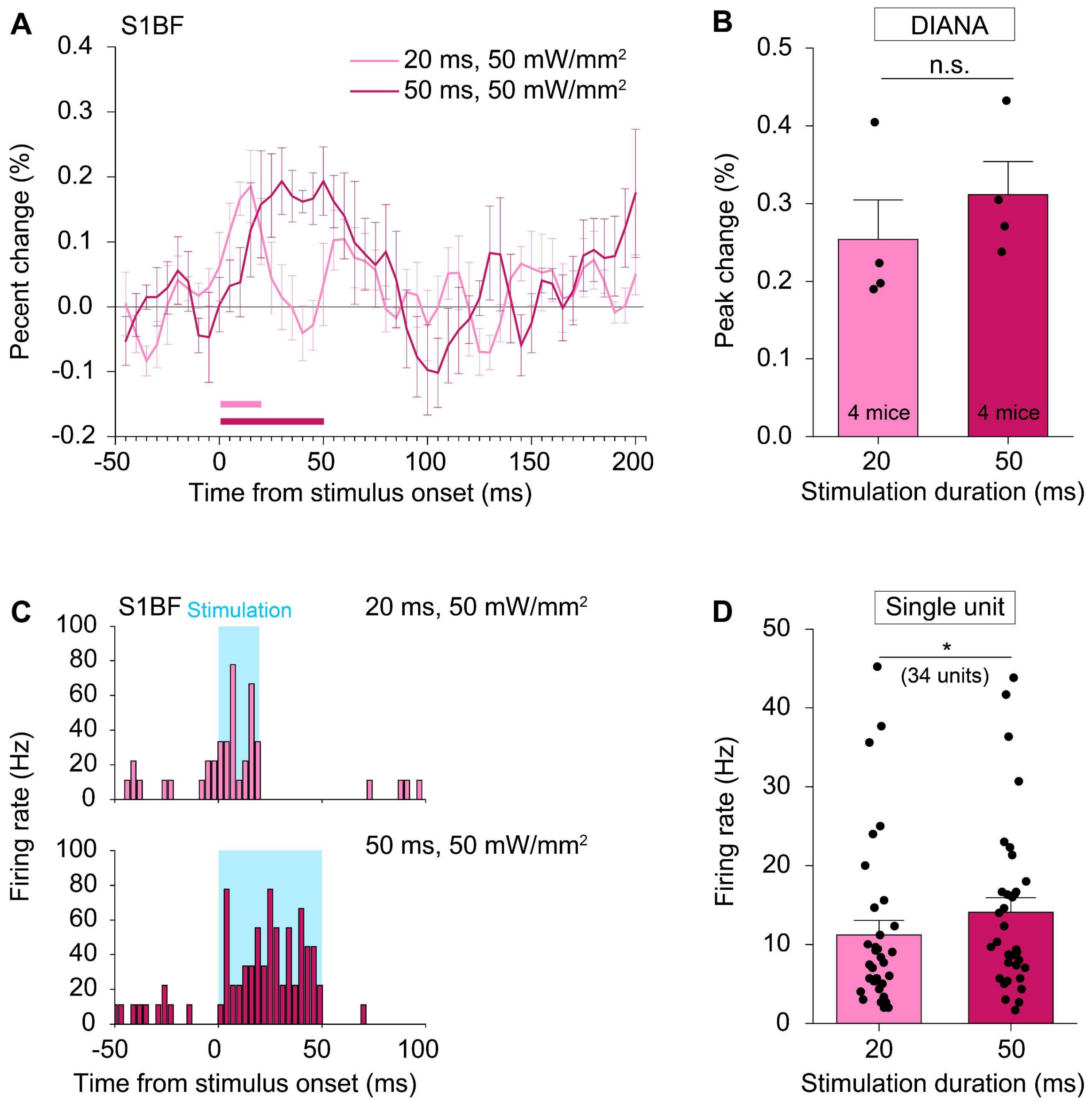


**Fig. S11. Dependence of DIANA response and spike firing rate on the duration of optogenetic stimulation.** (**A**) Optogenetic DIANA responses in the S1BF in response to 20 ms (light magenta) and 50 ms (magenta) blue light stimulation at the same intensity of 50 mW/mm$^2$ ($n$ = 4 mice). (**B**) Bar graph showing their peak percent signal changes. Horizontal bars in (A) indicate the duration of blue light stimulation at 20 ms (light magenta) and 50 ms (magenta). (**C**) Post-stimulation time histogram (PSTH) of the blue light stimulation-responsive single units over time in the S1BF responding to 20 ms (top) and 50 ms (bottom) blue light stimulation at the same intensity of 50 mW/mm$^2$. (**D**) Bar graph showing their spike firing rates ($n$ = 34 units from 8 mice). Blue shades in (C) indicate the durations of optogenetic stimulation, i.e., 20 ms (top) and 50 ms (bottom). All data are mean ± SEM. *: $p < 0.05$, n.s.: $p > 0.05$ for paired $t$-test.

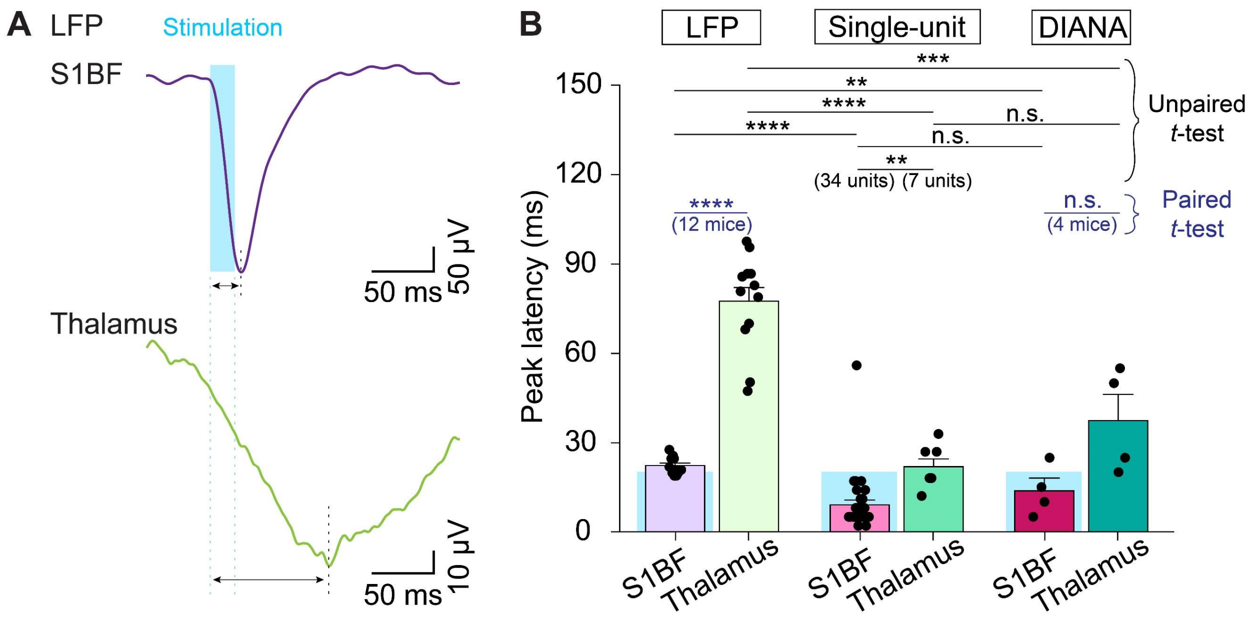


**Fig. S12. Comparison of DIANA response to local field potential (LFP) and single-unit spike in response to optogenetic stimulation of S1BF neurons *in vivo*.** (**A**) Representative LFP signals in the S1BF (top) and thalamus (bottom) evoked by blue light (473 nm) stimulation of 20 ms duration and 50 $mW/mm^2$ intensity. (**B**) Bar graph showing their peak latencies ($n$ = 12 mice) compared to the latencies of peak spike firing rates (S1BF, $n$ = 34 units from 8 mice; thalamus, $n$ = 7 units from 4 mice) and peak DIANA responses ($n$ = 4 mice). Blue shades/dotted lines in (A) and (B) indicate the duration of optogenetic stimulation in the S1BF. The vertical black dotted line in (A) indicates LFP peaks. All data are mean ± SEM. **: $p < 0.01$, ***: $p < 0.001$, ****: $p < 0.0001$, n.s.: $p > 0.05$ for paired and unpaired $t$-test.

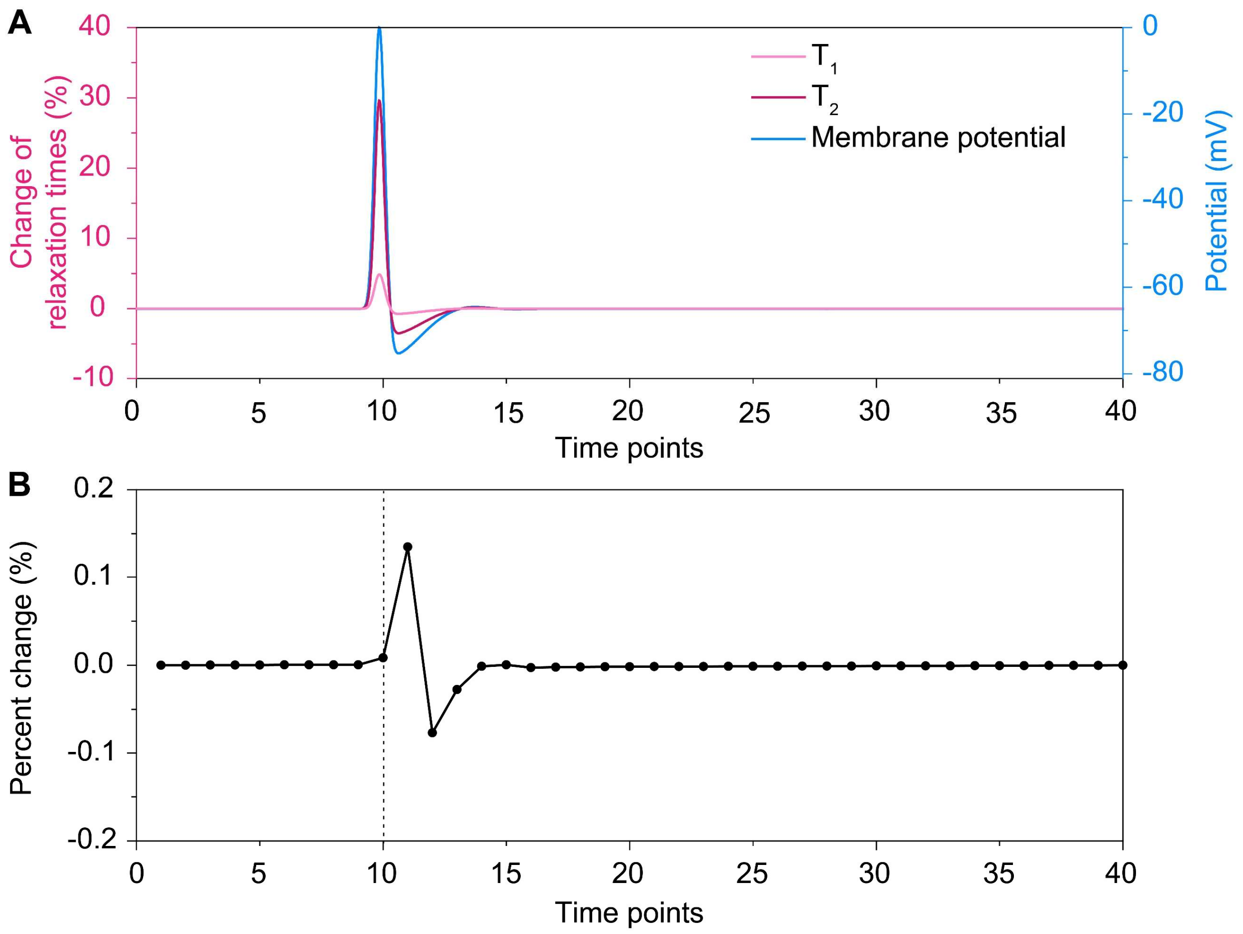


**Fig. S13. Simulation of DIANA signal change.** (**A**) Changes in $T_1$ (light magenta) and $T_2$ (magenta) relaxation times with varying membrane potential (blue), under the assumption of approximate, linear relationship between the relaxation times and membrane potential (**Fig. 4H**). (**B**) The DIANA signal change obtained using Bloch simulations based on $T_1$ and $T_2$ relaxation times given in (A). The interval between two consecutive time points was 5 ms.
